# Emergent Chemical Reactivity and Complexity of RNA Condensates

**DOI:** 10.1101/2025.04.15.648868

**Authors:** Samuel Hauf, Ryo Nakamura, Barbara Cellini, Yohei Yokobayashi, Mirco Dindo

## Abstract

The RNA world hypothesis posits the existence of life-like assemblies that consisted mostly of RNA. However, questions remain regarding the emergence of RNA catalysis, stability, reactant availability, and compartmentalization of genetic material. At acidic pH, short RNAs (average ≈ 20 nt) readily phase-separate into a condensed phase enriched with long RNA. These RNA condensates stably compartmentalize RNA as well as DNA and maintain stable identities even in the absence of membranes. Additionally, the RNA condensates concentrate ions, small molecules, phospholipids, peptides, ribozymes, and proteins. Beyond enriching such diverse components, RNA condensates function as microreactors with two catalytic capabilities: they physically enhance reaction rates by concentrating reactants within a confined space and simultaneously act as inherent catalysts that directly facilitate chemical transformations. RNA condensates can also support ribozyme and enzymatic activity. Together, these findings suggest that RNA phase separation may have played a crucial role in life’s origins by providing compartmentalization, inherent catalytic activity, and molecular enrichment of long, potentially catalytic biopolymers.

## Introduction

Many biomolecules — like nucleic acids, proteins, lipids, and cofactors — are necessary to sustain the complex biochemistry that powers living organisms. However, how these lifeless components initially assembled to form the first active entities remains one of science’s unanswered questions. The RNA world hypothesis^1^ offers a potential explanation by proposing that life began with assemblies composed entirely of RNA that functioned as both information carrier and catalyst.^2,3^ This hypothesis has gained significant traction in origin of life research due to the remarkable chemical versatility of RNA.^2,4–6^

Despite its explanatory power, the RNA world hypothesis faces several significant challenges.^7–9^ It does not explain how information-encoding or catalytically active RNAs originally emerged. Many prebiotic chemistries cannot efficiently catalyze the formation of long RNA polymers from monomers, because reactions in aqueous environments typically yield short oligomers a few nucleotides in length, ^10^ with polymer extension becoming increasingly improbable with each additional nucleotide — the “Flory Length Problem”.^11^ Long RNAs are difficult to obtain but can form from monomers on mineral surfaces^12,13^ or from short RNA oligomers through loop-closing reactions.^14^ Furthermore, RNA is often considered too complex and unstable for prebiotic environments,^7,15^ although recent evidence suggests that RNA might have been stable under acidic conditions of the early Hadean eon. ^16^

Another critical challenge is the catalytic efficiency of early biopolymers. Randomly polymerized RNA or protein catalysts would have initially existed at extremely low concentrations with minimal catalytic activity. Such unspecialized catalysts require high substrate and catalyst concentrations to function effectively.^7,9,15^

Compartmentalization provides a potential solution by concentrating key molecules and creating discrete units for natural selection.^17–20^ To explain how chemicals could be organized into life-like assemblies, various protocell models have been proposed.^21–27^ Compartments with membranes based on lipids or peptides are commonly considered plausible proto-cells,^28,29^ but they present significant limitations. Membrane impermeability to polar charged molecules creates barriers for protometabolism and information flow.^30,31^ In addition, such protocells require the simultaneous presence of at least two chemically distinct species (RNA plus lipids or peptides), which decreases the probability of spontaneous assembly in prebiotic environments. Micelles are another simple system that can exhibit functionalities, such as compartmentalization, enrichment, and catalysis. Some of the mechanisms contributing to micelle-mediated catalysis are: changes in the localization and concentration of reactants, changes in the chemical environment, such as hydrophobicity and charge, as well as changes in the stability of intermediates.^32^

Membrane-less compartments formed by liquid-liquid phase separation (LLPS) offer a potential alternative.^33,34^ In LLPS, a solution of previously miscible components spontaneously forms two distinct coexisting liquid phases: a dense phase enriched in specific components and a surrounding dilute phase depleted of these components. Progress has been made in producing membrane-less systems from various biomolecules, ^35–42^ with some demonstrating division,^38,43^ migration,^44^ and out-of-equilibrium behaviors.^45,46^ Although various membrane-less systems have been proposed, they typically require multiple distinct biopolymers or specific environmental conditions, making them unlikely in prebiotic settings. Recently, it has been suggested that the RNA condensates themselves could have been catalytically active, forming the units that drove early evolution at the origin of life.^47^

Our work experimentally explores how RNA alone can form membrane-less compartments through phase separation. Using commercially available yeast RNA preparations as a surrogate for short RNA sequences, we observed spontaneous RNA condensation under acidic conditions through phosphate backbone protonation. These RNA condensates exhibit several properties crucial to the origin of life. First, they selectively concentrate longer RNA species, indicating a way to circumvent the Flory Length Problem. Second, they stably compartmentalize RNA in the absence of a membrane, without observable exchange of compartmentalized nucleic acids between different condensate populations, even during long-term merging events. Third, a wide range of molecules partition into their interior. This partitioning allows substrates as well as catalysts to be enriched, addressing the efficiency problem of early catalysis.

We further demonstrate that the RNA condensates support non-enzymatic catalysis through distinct mechanisms. The enrichment of molecules within the condensates significantly enhances catalytic efficiency. Additionally, the RNA condensates act as inherent catalysts themselves, due to their intrinsic chemical properties. In addition to enhancing non-enzymatic biochemical reactions, the RNA condensates also support complex biocatalysis by ribozymes and multi-subunit enzymes, improving their potential as vessels for primitive metabolism. Intriguingly, RNA condensates spontaneously interact with short-chain fatty acids to form hydrophobic surface layers (representing a potential bridge to primordial membranes), while phospholipids introduced to these systems form circular sub-structures within the condensates, further increasing their molecular complexity.

RNA condensates could serve as nucleation points for increasing complexity, selectively enriching nucleotides, peptides, and lipids, allowing the emergence of more sophisticated functions. The encapsulation of diverse biomolecules within initially membrane-less proto-cells could have created the conditions necessary for primitive metabolism and replication, initiating cycles of mutation and selection that drove biological evolution. Our observations suggest a pathway linking prebiotic chemistry to the RNA world. This link of chemistry to biology by RNA phase separation suggests a potential pathway to address the challenges of concentration, compartmentalization, and catalysis at the origin of life.

## Results and Discussion

### RNA phase separation

RNA is an acid consisting of a hydrophilic sugar-phosphate backbone and hydrophobic nucleobases (Fig. 1 top). Concentrated nucleic acid solutions readily undergo phase separation upon dehydration.^48^ We show that in addition to dehydration, RNA solutions close to saturation easily phase-separate under a variety of conditions. Details on the RNA used can be found in the supplementary information (Section **”Characterization and identity of phase-separated RNA”** and Fig. S1). Storing concentrated RNA solutions (5-10 g/L) on ice causes them to become turbid with numerous condensate droplets visible under the microscope (Fig. 1, middle). The addition of multivalent cations, such as magnesium (Fig. 1, right and Fig. 2a)^49–52^ and others reported in Fig. S2, also triggers the phase separation of RNA in a concentration-dependent manner. We observed that a minimum of 25 mM MgCl_2_ is required to induce phase separation at RNA concentrations as low as 0.55 g/L (Fig. 2a, heatmap).

**Figure 1:**
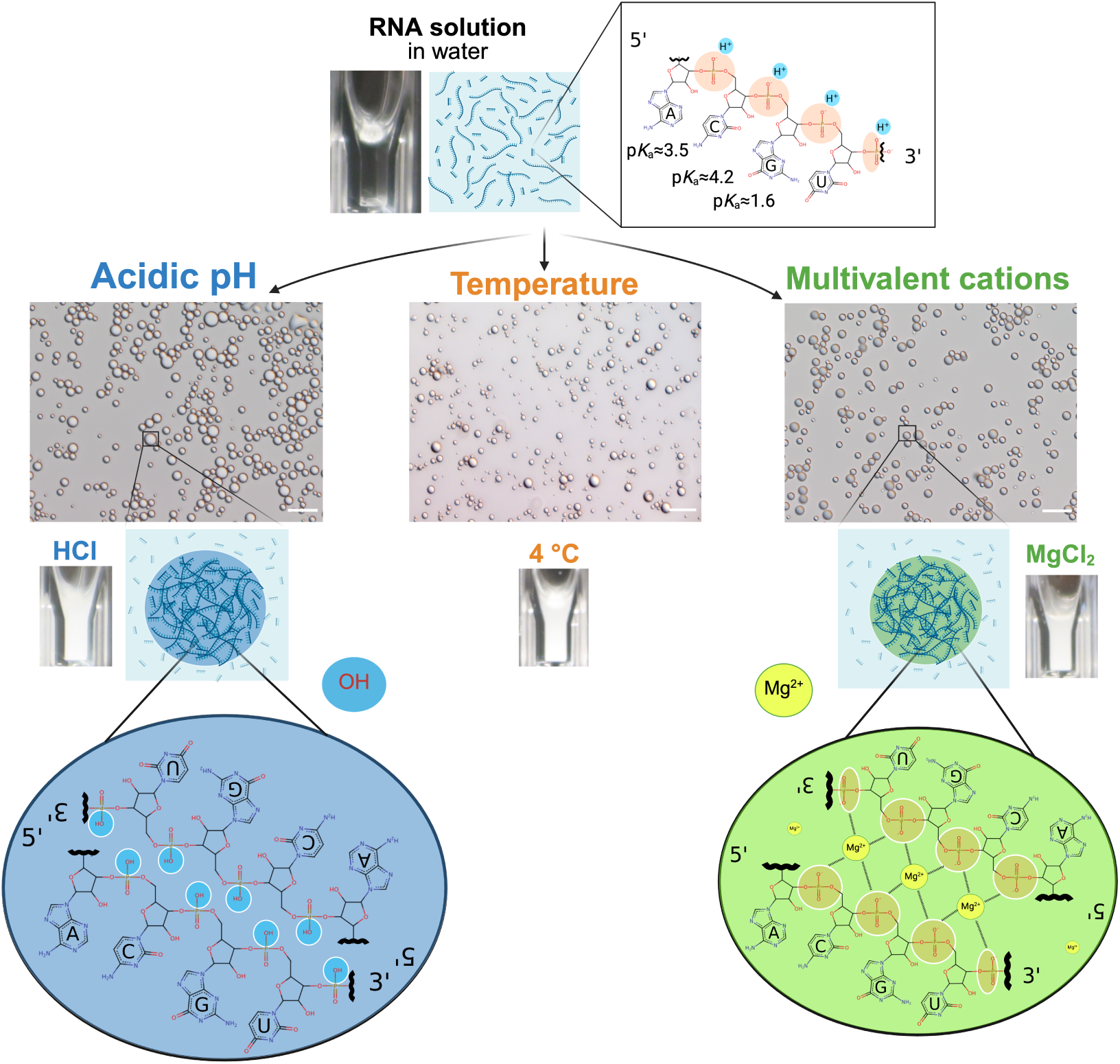
Overview of conditions inducing RNA phase separation from concentrated solu-tions. A near-saturated RNA solution (5-10 g/L at room temperature) forms phase-separated condensates under diverse conditions: decreasing pH with acids (left, blue), lowering tem-perature (middle), or introducing multivalent cations such as Mg^2+^ (right, green; see also Fig. S2). Microscope images show the resulting RNA condensates under each condition. Scale bars are 10 µm long. The molecular mechanisms promoting RNA condensation are illustrated for acidic conditions (protonation of phosphate groups, blue) and multivalent cations (electrostatic interactions with RNA strands, green), both leading to reduced RNA solubility by diminishing backbone charge repulsion.

**Figure 2:**
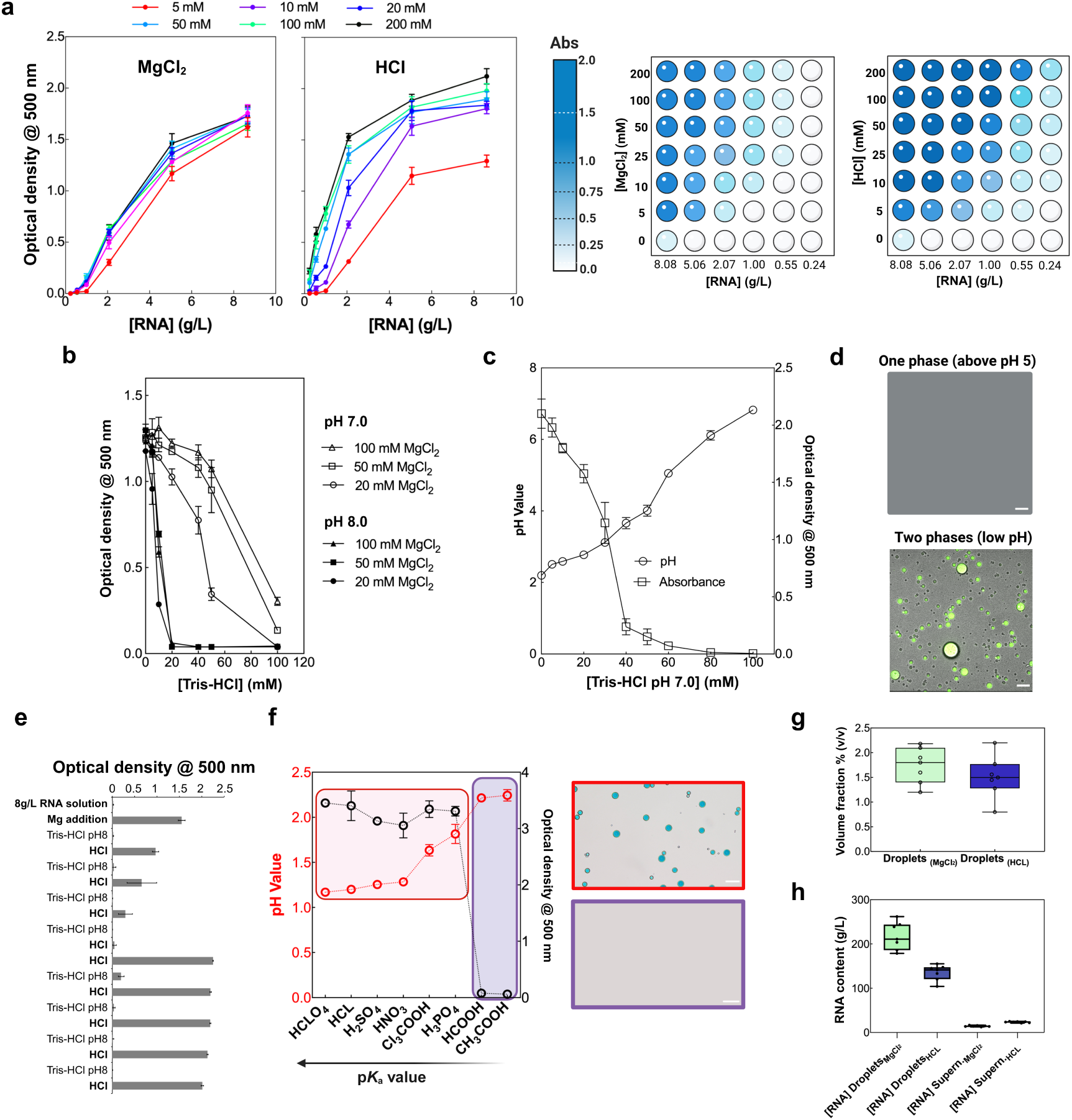
RNA phase separation mediated by divalent cations and by adding hydrochlo-ric acid. **a,** Optical density measurements showing RNA condensate formation conditions. Left: MgCl_2_-induced phase separation at different concentrations (5-200 mM) across vary-ing RNA concentrations (0-10 g/L). Right: HCl-induced phase separation under identical conditions. Heat maps display optical density values indicating condensate formation with darker blue representing higher turbidity. **b,** Turbidity decrease upon Tris-HCl (pH 7 and 8) addition at different concentrations (0-100 mM) for various MgCl_2_ concentrations. **c,** pH and turbidity correlation during Tris-HCl titration of condensates formed with 20 mM MgCl_2_, demonstrating the pH-dependent nature of phase separation reversal. **d,** Representative microscopy images showing the single phase (above pH 5, top) and two-phase regime (low pH, bottom with visible droplets). Green fluorescence represents FAM-signal of the labeled hammerhead ribozyme substrate partitioning into the RNA phase. Scale bars are 20 µm. **e,** Reversibility of RNA phase separation by buffer modulation. The addition of 10 µL of a 1 M magnesium chloride solution to solution of approximately 8 g/L RNA, 20 mM Tris-HCl pH 8 (460 µL) triggers phase separation, increasing optical density at 500 nm. Further addition of 15 µL 1 M Tris-HCl pH 8 causes the RNA condensates to dissolve due to the resulting increase in pH. Subsequent addition of 10 µL 0.5 M HCl causes the RNA condensates to re-form due to a drop in pH. This process was repeated for 8 cycles. HCl addition was not sufficient to fully compensate for the addition of Tris-HCl, so after 4 additions of Tris-HCl pH 8 two additions of HCl were necessary to induce phase separation again. This caused a more significant drop in pH, resulting in stronger phase separation. **f,** Correlation between acid strength (pK*_a_*) and phase separation efficiency. pH values (black circles) and optical density measurements (red circles) for different acids arranged by increas-ing pK*_a_* value. Strong acids (low pK*_a_*) effectively trigger phase separation while weak acids (high pK*_a_*) do not. Representative microscopy images of RNA solutions with added HCl (red box) and acetic acid (purple box). Scale bars are 20 µm. **g,** Quantitative comparison of droplet volume fraction and **h,** RNA content between MgCl_2_ and HCl conditions. Data presented are averages (*n* ≥ 3) with error bars indicating standard deviation. In the box plots, whiskers indicate the range.

Remarkably, RNA phase separation occurs even more readily when strong acids are added. Hydrochloric acid (pK*_a_* gas −5.9) is particularly effective (Fig. 1, left), inducing phase separation at concentrations as low as 10 mM even with minimal RNA concentrations (around 0.237 g/L; Fig. 2a). The critical role of pH in RNA phase separation is evident when manipulating buffer composition (Fig. 2b,c). Increasing the pH by titrating with Tris-HCl buffer (pH 7.0 or 8.0) dramatically reduces both the size and number of RNA condensates (Fig. S3), leading to complete dissolution above pH 4.5-5.0 (Fig. 2c,d). The process of condensate formation at low pH and dissolution at higher pH was repeated for 8 cycles, indicating the reversibility and robustness of RNA phase separation at low pH (Fig. 2e). RNA deamination was minimal under these conditions (Fig. S4), while alkaline hydrolysis of the RNA prevented phase separation (Fig. S5).

To further investigate the relationship between acidity and RNA phase separation, we tested acids with different pK*_a_* values (Fig. 2f). Strong acids such as nitric acid (pK*_a_* 1.4), or sulfuric acid (pK*_a_*_1_ = 2.8, pK*_a_*2 = 1.99) efficiently induce RNA phase separation, whereas weaker organic acids such as formic acid (pK*_a_* 3.745) and acetic acid (pK*_a_* 4.756) do not trigger RNA phase separation (Fig. 2f). By correlating pH measurements with absorbance data, we determined that protonation of the phosphate backbone (the pK*_a_*value of the backbone phosphodiesters is between 1 and 2^53,54^) plays a critical role in driving RNA phase separation. The solubility of RNA is primarily determined by the charge and hydrophilicity of its sugar-phosphate backbone, which is significantly affected by phosphate protonation. This explains why the solubility of RNA increases dramatically upon the addition of a base such as potassium hydroxide (Fig. S1a).

Our experimental data (Fig. 2), combined with the current literature, suggest a common mechanism underlying these diverse phase separation triggers. Low temperature reduces the solubility of RNA and influences RNA-solvent and RNA-RNA interactions.^55^ Strong acids protonate the phosphate backbone, reducing repulsive negative charges between RNA strands. Multivalent ions screen the backbone’s negative charges and potentially coordinate different RNA strands via electrostatic interactions ^56,57^ (Fig. 1, bottom). Thus, reduced RNA solubility (primarily through diminished backbone charge) allows attractive hydropho-bic interactions between the nucleobases to drive phase separation.

Since high ion concentrations (*>*20 mM MgCl_2_; Fig. 2a) may be unlikely in prebiotic environments with abundant RNA, we investigated whether organic co-solutes could enhance phase separation at low ion concentrations. We chose the organic co-solutes propylene glycol, 1,6-hexanediol, DMSO, and PEG for investigation. The first two are simple polyols of which at least propylene glycol might have come from space.^58,59^ DMSO and PEG were chosen as model molecular crowding agents. All of them significantly promote phase separation at low magnesium concentrations (10 mM; Fig. S1c). A relatively modest co-solute concentrations (5%) is enough to enhance phase separation, with increasing concentrations amplifying the effect (Fig. S1c).

RNA condensates produced with 200 mM HCl or MgCl_2_ had similar volume fractions (1-2%; Fig. 2g), but different RNA concentrations within the condensates. MgCl_2_-induced condensates contained higher RNA concentrations (approximately 200 g/L vs. 150 g/L for HCl-induced condensates; Fig. 2h), suggesting structural differences between these types of condensates.

The remarkable ease with which RNA phase-separates in response to diverse triggers — temperature, pH, multivalent cations, and molecular crowding — highlights its unique capacity for condensate formation, a property not readily observed with other macromolecules. Additionally, RNA condensates selectively enrich longer RNA species over shorter ones (Fig. S1b, Fig. S6, and Fig. S18), likely due to favorable enthalpic interactions from increased valency counterbalanced by lower entropic costs.^60,61^ The importance of RNA length is supported by the fact that increased shortening of the RNA by alkaline hydrolysis progressively reduces its capacity for phase separation (Fig. S5).

These properties facilitate phase separation and selective enrichment of longer RNAs, indicating a way to circumvent the Flory Length Problem in origin of life scenarios. The rare species of randomly produced longer RNA could be concentrated and compartmentalized within condensates. Assuming the catalytic capacity to polymerize nucleotides,^62^ new RNA molecules could have been synthesized within the compartments. This raises the question whether nucleotides and other relevant biomolecules could be enriched in these RNA condensates to enable catalytic activities.

### Partitioning of small molecules within RNA condensates

From an origin of life perspective, an important feature of a protocell is its ability to interact with and compartmentalize a wide variety of organic molecules coupled with its ability to catalyze chemical reactions. Guest molecules should have a variety of different physico-chemical characteristics and sizes to enable the complex chemistry essential to form entities capable of sustaining primordial chemical reactions. The enrichment of small molecules is therefore an essential property of any protocell system and has been reported for other systems.^36^

With these premises, we analyzed the partitioning of several organic guest molecules into RNA condensates as reported in Fig. 3 and checked the diffusion of small organic molecules within the condensates (Fig. S7). As expected, small organic molecules (such as rhodamine 6G) can easily enter the droplets and small aromatic fluorophores such as methylene blue, 4,6-diamidino-2-phenylindole (DAPI), resorufin, and SNARF are highly concentrated inside RNA condensates (Fig. S8). In addition, ATP was also enriched within RNA condensates, as was FMN (although to a lower extent) (Fig. 3).

**Figure 3:**
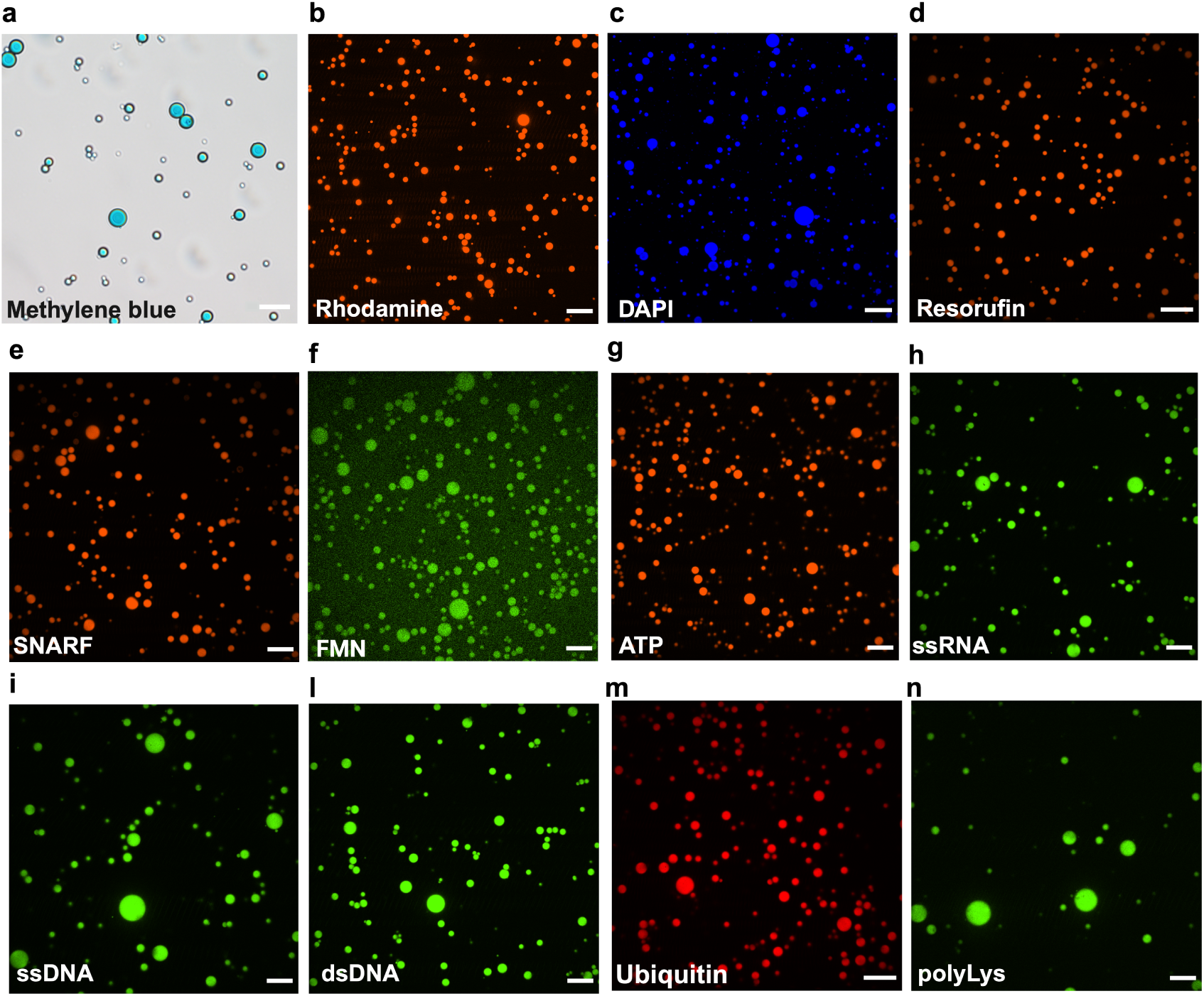
Partitioning of small organic molecules, nucleic acids and peptides within the RNA phase-separated condensates. **a,**RNA phase-separated condensates have been obtained by adding 10 mM HCl to 5 g/L RNA in the presence of **a,**Methylene blue. **b,**Rhodamine 6G. **b,** DAPI. **d,** Resorufin. **e,** SNARF™-4F 5-(and-6)-carboxylic acid. **f,** FMN. **g,** ATP visualized using BioTracker ATP-Red Live Cell Dye. **h,** FAM labeled ssRNA. **i,** FAM labeled ssDNA. **l,** FAM labeled dsDNA. **m,** Ubiquitin and **n,** FITC labeled polyLys. The concentration of small organic molecules partitioned in the RNA phase-separated condensates was between 200 nM and 20 µM. The same molecules at the same concentration can be partitioned by using 10 mM MgCl_2_ using 5 g/L RNA. The quantification of the signal within and outside the condensates is reported in Fig. S8.

Another important property of protocells is the encapsulation of longer, potentially information-coding biopolymers like nucleic acids. As shown in Fig. S1 the RNA condensates enrich longer species of RNA. RNA condensates prepared from solutions containing short RNA fragments using both MgCl_2_ and HCl (10 mM) are also enriched by longer labeled nucleic acids like RNA, single-, and double-stranded DNA (Fig. 3, Fig. S6, Fig. S18).

Several coacervate systems have the drawbacks that encapsulated molecules easily cross the membrane-less droplet boundary and that the coacervate phase quickly coalesces into one phase. The movement of encapsulated species across the boundary of membrane-less coacervate droplets and the coalescence of droplets are issues that have so far hampered their use as protocell models.^35,63,64^

The RNA system reported here limits this drawback. Even though the RNA condensates are liquid-like, showing fluorescence recovery after photobleaching (FRAP, Fig. S9), compartmentalization was remarkably stable. The mobility of RNA between the condensate droplets was tested by preparing three different RNA condensate populations with synthetic RNAs carrying different fluorescent labels (Fig. S6a). Although slow fusion events and wetting could be observed, the populations remained distinct for extended periods of time after mixing (Fig. S6b,c). This result demonstrates that RNA can be efficiently compartmentalized within RNA condensates, maintaining (during the limited experimental time) separate genetic identities. Phase-separated RNA thus seems to fulfill some essential requirements for a protocell system^65^ that have been difficult to achieve with other phase-separated systems, such as reduced Ostwald ripening and efficient compartmentalization of RNA species required for Darwinian-style evolution.

An important role of RNA in extant biochemistry is the assembly of peptides from amino acids, which supposedly is a remnant of the RNA world.^66^ Therefore, we investigated how the RNA phase interacts with amino acids and peptides. Interestingly, RNA condensates partition peptides and small proteins of different length. As reported in Fig. 3, ubiquitin (8.6 KDa, 76 aa) and the longer poly-Lys (15-30 KDa) are enriched within the RNA phase while amino acids are roughly equally distributed between both phases (Fig. S10). Besides interacting with small molecules and peptides, the RNA condensates also partitioned phospholipids, which spontaneously formed sub-structures inside the RNA condensate. Fatty acids (C11 and C12) spontaneously assembled on the condensate surface (Fig. S11) that resemble a primordial membrane layer.

In summary, many different small molecules and even more complex biopolymers partition into RNA condensates. Importantly, longer RNA species and nucleotides are enriched. The enrichment of molecules and the ease of RNA phase separation, without the need for other polymers or components, are promising indications for the potential of RNA conden-sation in the context of the origin of life. The findings also open the door for investigations into the catalytic potential of RNA condensates as microreactors for chemical reactions.

### RNA condensates as prebiotic microreactors: catalytic enhancement of simple chemical reactions

Driven by the enrichment of molecules within RNA condensates, we investigated whether these compartments could catalyze or enhance simple chemical reactions relevant to prebiotic chemistry. Hydrolysis, condensation, and transamination reactions would have been critical for the formation and interconversion of the building blocks necessary for life. Hydrolysis is fundamental to metabolism and energy transfer in living systems,^38^ while condensations allow the formation of complex molecules from simpler precursors. Transamination, the transfer of amino groups between molecules, represents a key process for amino acid synthesis and nitrogen cycling in primitive metabolic networks.^67,68^

We first examined the hydrolysis of para-nitrophenyl acetate (pNPA, Fig. 4a), a widely used substrate for evaluating esterase activity that has been previously employed to study catalysis in compartmentalized systems.^69,70^ The reaction can be followed by monitoring pNP absorbance (Fig. 4b). Under our experimental conditions (pH 2.0 −2.5), the acid-catalyzed hydrolysis of pNPA is extremely unfavorable: the mechanism requires protonation of the ester carbonyl and water is a particularly weak nucleophile in acidic environments. Our kinetic analysis revealed reaction rates of 3.0 × 10^−9^ M/s for 4 mM pNPA in RNA condensates, compared to 1.9 × 10^−9^ M/s in the diluted phase. Statistical analysis (t-test) showed a significant difference between condensates and the diluted phase (p ¡ 0.05), but no significant difference between condensates and water alone (p ¿ 0.05). Although this enhancement is modest, the fact that we observed a trend toward acceleration under such unfavorable conditions suggests that RNA condensates may provide alternative reaction pathways, potentially through specific functional groups within the RNA structure serving as general acid-base catalysts or through creating microenvironments with altered local reactivity compared to bulk solution.

**Figure 4:**
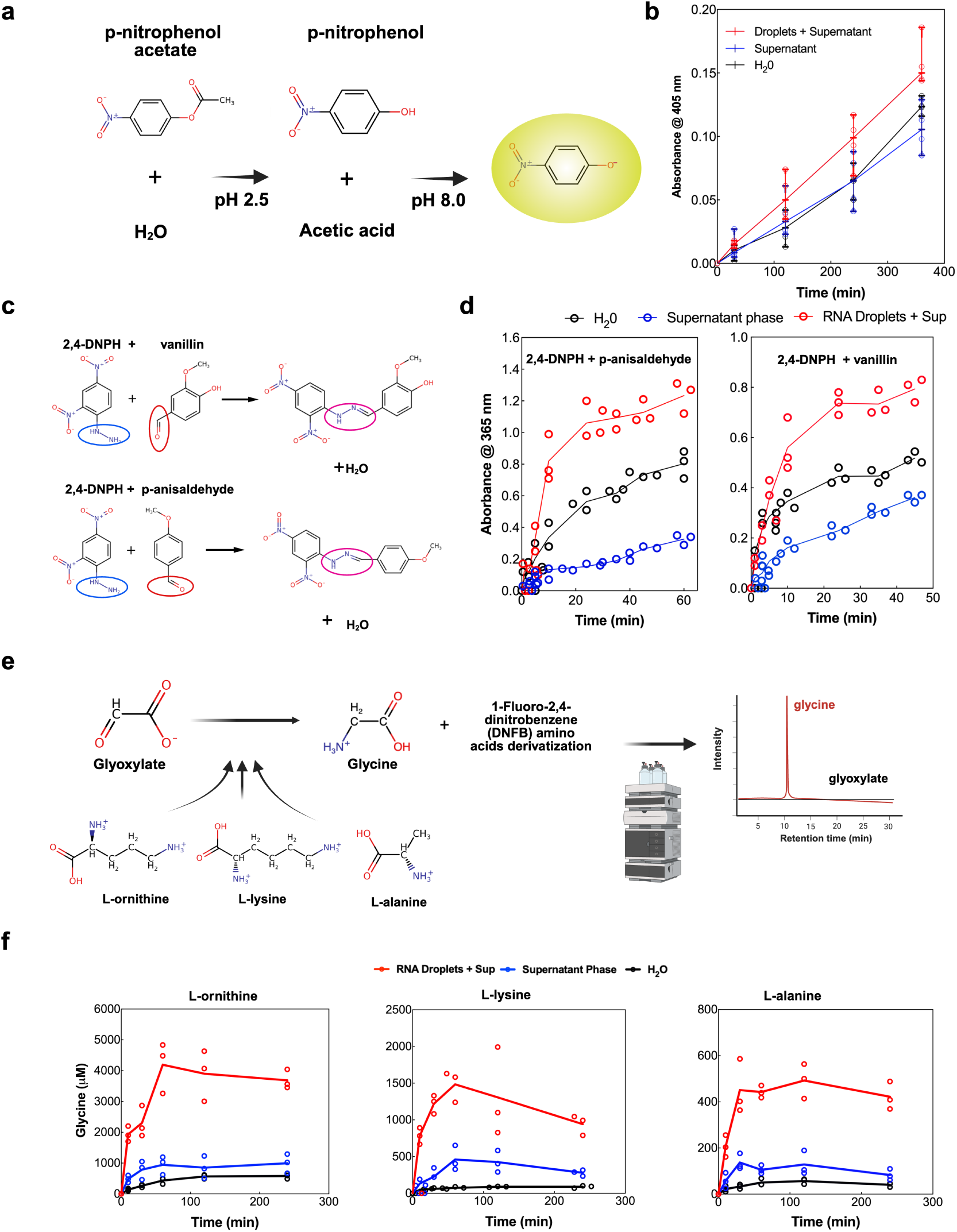
Small organic molecule catalysis within the phase-separated RNA condensates. **a,** Chemical scheme showing enhanced acid hydrolysis of p-nitrophenol-acetate (pNPA) reaction mechanism. **b,** Time course analysis of 4 mM pNPA hydrolysis in RNA condensates (red dots and line) compared to the supernatant phase (blue dots and line) and control reaction in water alone (black dots and line) in the presence of 100 mM MgCl_2_. The reaction produces p-nitrophenol (yellow at neutral and basic pH) and acetate. Data presented are averages (*n* ≥ 3) with error bars indicating standard deviation. Fig.S16 shows additional pNPA hydrolysis experiments at 1 mM concentration as well as the experimental procedure schema. **c,** Chemical schemes for addition reactions between 2,4-dinitrophenylhydrazine (2,4-DNPH) and aldehydes: conversion of p-anisaldehyde to p-anisaldehyde,2,4-dinitrophenylhydrazone (top) and vanillin to vanillin,2,4-dinitrophenylhydrazone (bottom). **d,** Reaction kinetics showing formation rates of the hydrazone products. 2,4-dinitrophenylhydrazine was used at a concentration of 200 µM and the aldehydes at 400 µM, with 100 mM MgCl_2_ to trigger phase separation. Left panel: p-anisaldehyde reaction; Right panel: vanillin reaction. Fig.S12 contains raw data and complete absorbance graphs. **e,** Chemical scheme illustrating glycine production from transamination reactions using different amino donors (L-ornithine, L-lysine, L-alanine) with glyoxylate as the *α*-ketoacid acceptor, followed by HPLC analysis workflow. **f,** Time course of glycine production within RNA condensates (red dots and line) compared to the supernatant phase (blue) and control in water (black). Experimental conditions: amino donors (L-ornithine, L-lysine and L-alanine) at 10 mM, glyoxylate at 20 mM. Following incubation and derivatization with 1-fluoro-2,4-dinitrobenzene (see Materials and Methods), amino acids were quantified using HPLC. Fig. S10 reports HPLC chromatograms and amino acid calibration curves. Control experiments in water maintained identical MgCl_2_ concentrations (100 mM) and acidic pH (adjusted with HCl) as the condensate samples.

More significant effects were observed with the formation of 2,4-dinitrophenylhydrazones (Fig. 4c, d), a model condensation reaction. Condensation reactions are crucial in prebiotic chemistry for the formation of carbon-carbon and carbon-nitrogen bonds to build complex organic molecules from simple precursors. The reaction of 2,4-dinitrophenylhydrazine (2,4-DNPH, 200 µM) with vanillin or p-anisaldehyde (400 µM) proceeded substantially faster in RNA condensates compared to control conditions. For p-anisaldehyde, we measured a rate constant of 4.5 ± 0.6 · 10^−4^µM^−1^min^−1^ in RNA condensates, compared to 5.0 ± 0.5 · 10^−5^µM^−1^min^−1^ in the supernatant phase, representing an 9-fold acceleration. Similarly, for vanillin, the rate constant increased from 2.6±0.3·10^−5^µM^−1^min^−1^ in the supernatant to 1.7± 0.3 · 10^−4^µM^−1^min^−1^ in RNA condensates, a 6.5-fold enhancement. The improved reactivity correlates closely with the observed 6-7 fold higher concentration of 2,4-DNPH within the condensates (Fig. S12d), providing quantitative evidence that molecular partitioning is the primary driver of the catalytic effect in this case (concentration effect).

Perhaps most remarkably, RNA condensates showed a robust enhancement of transamination reactions (Fig. 4e,f). Glyoxylate was converted to glycine at acidic pH using various amino acids as donors. Glyoxylate is considered an abiotically accessible organic compound and important prebiotic building block linked to protometabolism, ^71–73^ while glycine, the simplest amino acid, is fundamental for early peptide formation^74^ and has been detected in meteorites and prebiotic simulation experiments.^75^ The reaction proceeded efficiently at room temperature in the presence of RNA condensates, but less so in the supernatant phase or water (Fig. 4f). Using L-ornithine as the amino donor, the glycine concentration reached approximately 4192 µM in the presence of RNA condensates, compared to only about 995 µM in the supernatant phase and 483 µM in water. The reaction rates showed even more dramatic differences: 107 µM*/*min in the presence of RNA condensates versus just 14 µM*/*min in the supernatant and 10 µM*/*min in water. Similar enhancements were observed using L-lysine (38 µM*/*min in condensates vs. 2 µM*/*min in water) and L-alanine (14 µM*/*min vs. 1 µM*/*min in water) as amino donors.

This transamination reaction is typically unfavorable at acidic pH because the protonation of amino groups reduces their nucleophilicity and impedes the formation of imine intermediate essential for the reaction mechanism (see SI for detailed discussion). Despite these challenging conditions, RNA condensates dramatically enhance the amination of glyoxylate. In fact, the reaction did not proceed efficiently in water or in the diluted phase. In a setup with high RNA concentrations but without triggering phase separation (without condensate formation by addition of MgCl_2_ 100 mM) the reaction was also inefficient (data not shown). Importantly, transamination also occurred between glycine, probably the most abundant amino acid under prebiotic conditions, as the donor and pyruvate (Fig. S13). This demonstrates that RNA condensates function as efficient microreactors for such transamination reactions by combining selective enrichment with intrinsic catalytic capabilities. The synergistic effects of RNA structure, magnesium ions, and amino acid properties create an environment that significantly enhances reaction rates and yields, particularly under conditions that would otherwise be unfavorable. These findings provide valuable insights into potential mechanisms of prebiotic chemical evolution and highlight the remarkable catalytic potential of RNA-based compartmentalization.

### Folding and catalytic activity of macromolecules within RNA condensates

We have shown that RNA solutions easily phase-separate under a variety of conditions and that the phase-separated RNA condensates enrich longer RNA species and different small organic molecules, thereby enhancing different organic reactions such as hydrolysis, addition reactions, and transamination. An additional challenge for these condensates to be a viable protocell system is their capacity to support specialized catalytic activity of biopolymers.

On first sight, the conditions inside the condensed RNA phase seem unfavorable for biopolymer catalysis. Condensates form at high RNA concentrations and low pH. At high nucleic acid concentrations, non-specific base pairing could disrupt proper folding of ribozymes. Essential ions (such as magnesium) could also be sequestered by other RNAs. Further, most enzymes and ribozymes are not active at pH below 4 in buffer; however, RNA condensates could provide an altered chemical environment by different mechanisms, such as local buffering, altered solvation, or modified pK*_a_* values. This assumption of a condensate-specific chemical environment is supported by measurements using LysoSensor, indicating that the condensates were more acidic than the supernatant (Fig. S14). Additionally, when tracking resorufin *β*-D-galactopyranoside hydrolysis, the reaction seems to occur or start at the condensate boundary (Fig. S15). Therefore, experimental proof of biologically relevant catalytic activity of ribozymes and protein enzymes inside the condensed RNA phase is necessary.

Fluorescently labeled BSA and GFP could be observed within the RNA condensates, indicating partition of the proteins into the condensates (Fig. 5a,b). Furthermore, GFP fluorescence data indicate that protein folding is maintained inside the RNA condensates, because GFP loses its fluorescence upon unfolding. ^77^ On the basis of the protein partitioning data, we decided to evaluate if protein catalysis was possible inside the condensates. For this purpose, we used Oxalate decarboxylase (OxDC) from *Bacillus subtilis*. OxDC is a homohexamer (a dimer of trimers) of 264 kDa which belongs to the cupin superfamily and requires Mn^2+^ and O_2_ to catalyze the conversion of oxalate to formate and CO_2_. Notably, the pH optimum of this enzyme is at pH 4.2 but with a wide range of activity. ^78^ As reported in Fig. 5c,d, OxDC partitioned into RNA condensates and also showed activity inside. Although the enzymatic activity measured within the condensates (11 ± 2 s^−1^) is low compared to that obtained in diluted conditions in buffer at pH 4.2 (74 ± 6 s^−1^) using 10 mM oxalate,^76^ it confirms that enzymes can be active inside RNA condensates. Notably, the product of the enzymatic reaction (formate) dissolves the droplets (Fig. 5b and Fig. S17).

**Figure 5:**
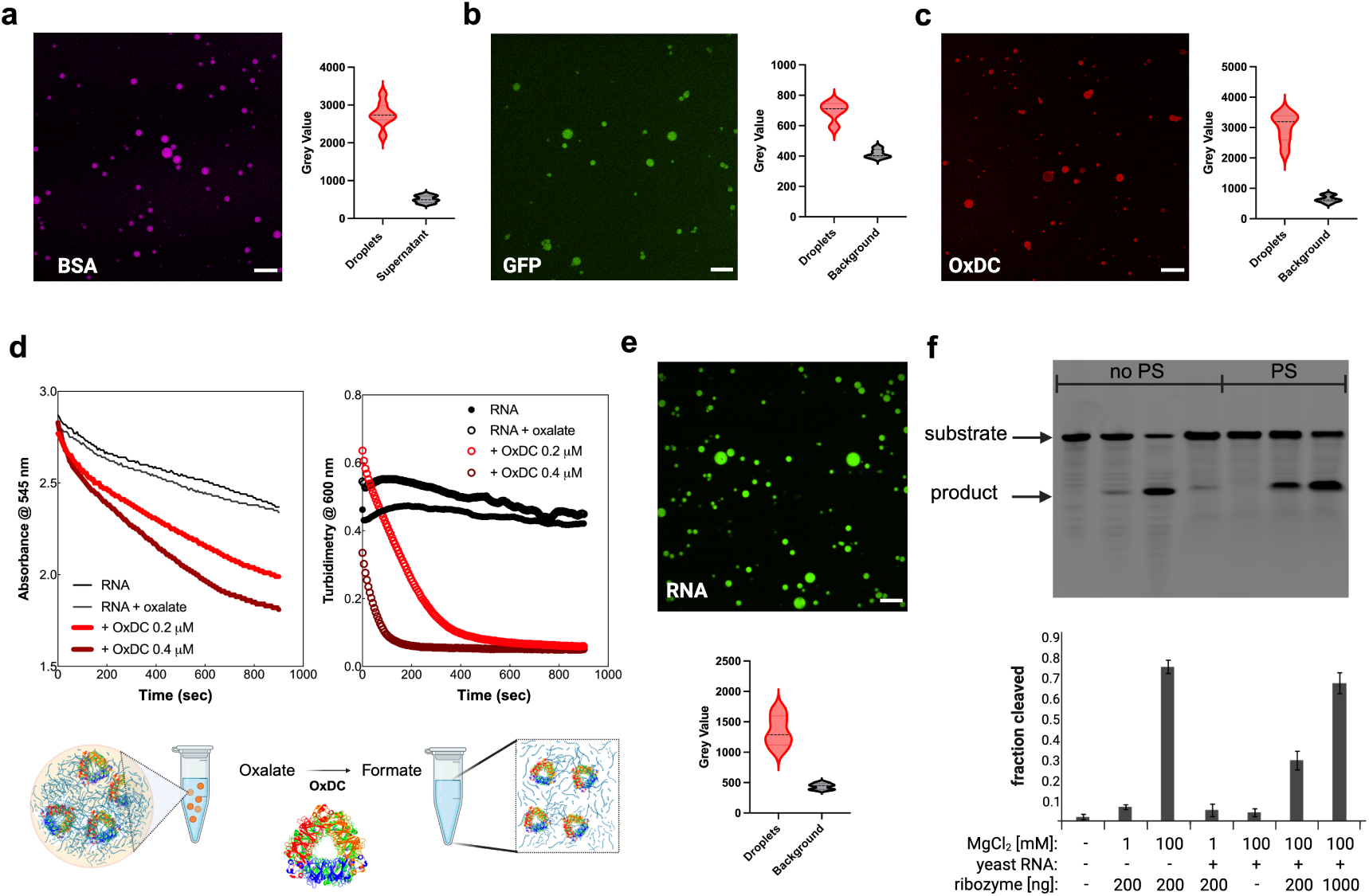
RNA condensates can accommodate catalysis mediated by macromolecules. Demonstration of protein partitioning and enzymatic activity within RNA phase-separated condensates. Protein partitioning experiments: **a,** 10 µM BSA labeled with Alexa Fluor 647 nm, **b,** 20 µM GFP and **c,** 5 µM OxDC partition efficiently within the condensates prepared using Tris-HCL 32 mM pH 7.5 with 3-3.5 g/L RNA and 100 mM MgCl_2_. Proteins were added immediately before MgCl_2_ addition to trigger phase separation. The fluorescence intensity analysis (grey-scale value) quantifies the signal enrichment within condensates compared to the surrounding solution. **d,** OxDC enzymatic activity within RNA condensates.^76^ Left: OxDC activity kinetics (light and dark red lines) measured using two enzyme concentrations: 0.2 µM and 0.4 µM in the presence of 10 mM oxalate substrate. Black lines show control experiments: RNA alone and RNA with oxalate substrate (10 mM final concentration) demonstrating background signal changes. Right: Turbidimetry measurements at 600 nm recorded during OxDC catalysis. The decreasing signal at 600 nm corresponds to RNA condensate dissolution caused by formate production (product of the oxalate decarboxylation reaction). **e,** Fluorescence microscopy showing FAM-labeled ribozyme substrate localization inside RNA condensates and quantitative analysis of the fluorescence signal enrichment in condensates compared to background. **f,** Top: Representative polyacrylamide gel electrophoresis (PAGE) analysis. Bands correspond to the FAM-labeled ribozyme substrate and the FAM-labeled cleavage product of the ribozyme reaction. Bottom: Quantification of substrate cleavage efficiency as averages with standard deviation derived from gel densitometry analysis of at least three independent experiments. Experimental conditions with and without phase separation (PS) are indicated. Lane-by-lane analysis: Lane 1: substrate remains stable in buffer alone. Lane 2: ribozyme activity at pH ≈ 3 with 1 mM MgCl_2_ shows weak cleavage (around 10%). Lane 3: increased MgCl_2_ concentration (100 mM) enhances cleavage to around 75%. Lane 4: addition of 5 g/L yeast RNA with 1 mM MgCl_2_ does not trigger phase separation and maintains cleavage efficiency similar to lane 2. Lane 5: phase separation induced by 100 mM MgCl_2_ does not affect substrate stability in the absence of ribozyme. Lane 6: ribozyme activity within phase-separated condensates shows reduced cleavage (around 30%) compared to bulk solution. Lane 7: increasing ribozyme concentration 5-fold within condensates achieves nearly 70% substrate cleavage, demonstrating dose-dependent activity.

Although the results obtained with proteins are promising, for the system to be a relevant protocell model inside the RNA world framework, it needs to be able to support nucleic acid-based catalysis. Therefore, the catalytic activity of nucleic acids was investigated using a hammerhead ribozyme.^79^ The ribozyme partitioned well into the RNA phase as shown on denaturing gels after separating condensates and supernatant by centrifugation (Fig. S18). The hammerhead ribozyme showed activity in RNA condensates formed from yeast RNA at 100 mM MgCl_2_ (Fig. 5e, f) or when adding PEG8000 at high concentrations (around 20%; data not shown). Although the activity of the ribozyme was less than 50% compared to the dilute condition without yeast RNA, the catalytic activity of the hammerhead ribozyme obtained inside the RNA condensates is promising.

This ribozyme is not adapted to the conditions used (low pH and high RNA crowding). Despite the many issues that could interfere with ribozyme catalysis, the activity observed was robust and dependent on the ribozyme concentration (Fig. 5f). A 5-fold increase in ribozyme concentration increased substrate cleavage to a level similar to that observed in dilute conditions. This shows that ribozymes can function inside the RNA condensates and we propose that in vitro selection of ribozymes under these conditions could yield ribozymes with good activity. The high RNA concentrations inside the condensates (*>*100 g/L) together with their catalytic capabilities and the enrichment of other molecules might be particularly favorable for the evolution of ligase or replicase ribozymes that could start evolutionary processes in the RNA world.

## Conclusion

RNA might have played a pivotal role in the emergence of life by performing multiple critical functions: storing genetic information, catalyzing biochemical reactions, and acting as a template for replication. Our findings demonstrate that RNA readily phase-separates under acidic conditions without requiring other biopolymers or small molecules. However, the resulting RNA condensates selectively enrich small molecules, peptides and more complex macromolecules such as functional ribozymes and enzymes.

Importantly, RNA condensates enhance biochemical reactions. They do so through two different mechanisms: First, through a concentration effect, these condensates create micro-environments where substrates and potential reaction partners are brought into close proximity, significantly increasing the probability of productive molecular interactions (addition reaction). Second, through an inherent catalytic effect, the RNA itself possesses intrinsic catalytic properties that can accelerate specific biochemical reactions independently of the concentration advantages (transamination reaction). These two distinct effects — local concentration of substrates and intrinsic catalytic properties — enable RNA condensates to enhance fundamental biochemical reactions even in the absence of dedicated catalysts. The RNA condensates can potentially facilitate other reactions that were not investigated.

Although our system does not directly address the abiotic origins of RNA, it provides substantive support for the possibility of the emergence and evolution of an early RNA world under acidic conditions. Geological evidence suggests that early Earth’s surface environments could have been moderately acidic, providing favorable conditions for RNA phase separation.^16^ This scenario would address the issue of RNA stability, supporting the hypothesis that the RNA world evolved at acidic pH. ^80^ Acidic conditions further facilitate RNA-RNA strand separation,^81^ essential for replication processes. A legitimate concern relates to RNA functionality under acidic conditions; however, hammerhead ribozyme activity has been demonstrated at pH values as low as 3,^82^ indicating that low pH does not necessarily impede RNA folding and function. Furthermore, RNA phase separation can be triggered by various metal ions that would likely be sequestered within RNA condensates where they could enhance catalysis.

The stable encapsulation of genetic material within the condensates indicates a plausible mechanism for maintaining genetic information within discrete catalytically active compartments. This is an advantage over previous phase-separated systems that easily coalesce, because it allows evolution of stable hereditary units into more complex protocells. Such compartmentalization could potentially have facilitated the gradual accumulation of active RNA sequences, thereby contributing to the molecular selection of condensates with enhanced functionality or stability. As such systems evolved, they could have developed and profited from interactions with other environmentally available molecules, for example fatty acids,^83^ which could have driven the development of increasingly complex boundary structures. These interactions may represent a coherent pathway toward the eventual emergence of membrane-bound protocells from initially membrane-free RNA condensates.

## Supporting information

RNA ICP data

## Acknowledgements

The research was supported by the Okinawa Institute of Science and Technology Graduate University (OIST) with subsidy funding to Y.Y.. M.D. thanks the University of Perugia for the support and the Japan Society for the Promotion of Science (JSPS) for the Kakenhi Early Career Scientist N. 22K15065, and R.N. acknowledges support from MEXT’s ‘Initiative for Realizing Diversity in the Research Environment. We are grateful for the help provided by the imaging section of the Core Facilities at OIST; in particular we thank Dr. Paolo Barzaghi for the great support with the confocal microscopy. The authors thank Prof. Claudio Constantini, Prof. Davide Chiasserini, Prof. Paola Laurino, Dr. Alessandro Bevilacqua and Dr. Franco Cardone for critical reading. The authors thank Yoshiteru Iinuma from the Instrumental Analysis Support at OIST for performing the trace element analysis. The images were prepared using *biorender.com*. We apologize to all the researchers involved in the field who have not been cited in our manuscript for space limitation.

