## Supplemental information for "Emergent Chemical Reactivity and Complexity of RNA Condensates"

### Supplementary Information for "Emergent Chemical Reactivity and Complexity of RNA Condensates"

### Experimental

#### Materials

RNA from yeast was from Roche (10109223001) or Fujifilm-Wako (4987481380561). RNA from torula yeast was purchased from Merck (R6625). NTP stock solutions (100 mM) were obtained from NEB. RNAse free water, propylene glycol, 1,6-hexanediol, dimethylsulfoxide (DMSO), and polyethylene glycol 200 (PEG200), HCl, sodium chloride, urea were purchased from Nacalai Tesque. Tris-HCl (varying pH from 7 to 9) and $MgCl_2$ solutions were ordered as 1 M stock solutions from Nippon Gene. The different salts tested were calcium chloride (Fujifilm Wako Pure), manganese sulfate (33341 Alfa Aesar), iron II sulfate (14498 Alfa Aesar), cobalt II chloride (255599 Merck), zinc chloride (208086 Merck), copper chloride (751944

Merck), sodium molybdate (243655 Merck), iron III chloride (F2877 Honeywell). PEG 200 was purchased from Tokyo Chemical Industry (TCI). L-Amino acids (L-alanine (A7627), L-ornithine (O2375), L-lysine (L5501), L-norvaline (D7627) and glycine (G7126)), DNFB (D1529), 2,4-dinitrophenylhydrazine (DNPH, D199303), p-anisaldehyde (A88107), vanillin (V1104), glyoxylate sodium salt (G4502), pNPA (N8130), pNP (241326), Rhodamine-6G (R4127), resorufin sodium salt (R3257) methylene blue (M9140), poly-L-lysine FITC labeled (P3543), DAPI (D9542), bovine serum albumin (BSA, A7638), $\beta$-Nicotinamide adenine dinucleotide, reduced disodium salt hydrate (NADH) (N8129), Pyruvate (P8524), Pyridoxal 5'-Phosphate (PLP), rabbit muscle L-lactate dehydrogenase and isopropyl-$\beta$-D-thiogalactoside were purchased from Merck (Germany). Fatty acids (D3835, D3821 and D3861), phospholipids ($\beta$-BODIPY™ FL C12-HPC (2-(4,4-difluoro- 5,7-dimethyl- 4-bora- 3a,4a-diaza-s-indacene- 3-dodecanoyl)- 1-hexadecanoyl-sn-glycero- 3-phosphocholine)(D3792), SNARF™-4F 5-(and-6)-carboxylic acid (C1272), acetoxymethyl ester, acetate (S23921), Alexa Fluor 647 Microscale Protein Labeling Kit (A30009), and LysoSensor™ Yellow/Blue DND-160 (L7545), were purchased from Thermo Fisher Scientific.

#### BSA, OxDC and ubiquitin labeling

BSA, OxDC and uniquitin from bovin liver erythrocytes were tagged using Alexa Fluor 647 Microscale Protein Labeling Kit, using the manufacturer's protocol but repeating the dialysis step three times to remove the free dye in the protein solution as reported.[1]

#### Preparation of RNA solutions

RNA solution was prepared by adding 200 mg of yeast RNA in 15 mL of DNAse/RNAse free water in 15 mL centrifuge tubes. The mixture was heated to 50 °C and vigorously shaken to dissolve as much RNA as possible. Subsequently, the solution was cooled to room temperature for at least 30 minutes so that the RNA partially precipitated. The saturated mixture obtained was then centrifuged at 10,000 g for 10 minutes at room temperature.

The supernatant was transferred to a new tube and used immediately or frozen at -20 °C. The frozen stocks were thawed at 50 °C and vigorously shaken from time to time until all precipitate was dissolved; thawed stock solution was cooled to room temperature before use. RNA concentrations were quantified by NanoDrop or by using MultiSkan SkyHigh Microplate reader (Thermo Fisher) using the microdrop Duo Plate. The RNA solution after resuspension in water has been diluted and read at 260 nm in triplicate every time and before use.

**Denaturing PAGE**

Denaturing PAGE was performed following standard procedure using 8 M urea PAGE gels. Shortly, 15% acrylamide gels containing 8 M urea were prepared from 40% pre-mixed stock of acrylamide/bis-acrylamide solution (40% acrylamide/Bis Mixed Solution (19:1), Nacalai 06140-45). RNA samples were loaded after resuspension and heat denaturation (2 min, 95 °C, then storage on ice) in TBE-Urea Sample Buffer / RNA loading dye (Thermo Fisher Scientific, LC6876) and run in TBE at 250 V for 50 minutes. Labeled RNA was detected directly while unlabeled RNA was stained with 1x SybrGold (Thermo Fisher Scientific, S11494) in 40 ml 1x TBE. Pictures were acquired using the Typhoon FLA 7000 (GE).

**RNA phase separation**

RNA phase separation has been triggered using RNA solutions prepared as described, both using stock solution or by diluting the RNA stock solution in water on in Tris-HCl buffer. For example, storing the solutions at lower temperature (for example around 0 °C on ice), adding different salts (such as $MgCl_2$, $CaCl_2$, $MnSO_4$, $ZnCl_2$, $CoCl_2$, $FeSO_4$, $FeCl_3$, $Na_2MoO_4$ prepared as 1 M stock solutions in water), strong acids (hydrochloric acid, sulfuric acid, trichloroacetic acid and perchloric acid) caused RNA to phase-separate (used at different concentrations). The exact conditions are described in the manuscript. To estimate the volume fraction, condensates from a total of 10 mL were left to settle over night in 15 mL centrifuge tubes. The tubes were then flicked carefully and left for another 30 minutes to allow condensates from the side of the tube to settle to the bottom. The supernatant was carefully removed before the volume of the condensate phase was estimated by pipette. For the estimation of RNA content, the condensates were resuspended in 15 mL water with heating to 50 °C before measuring with absorption at 260 nm. Resuspension as well as volume estimation values provided are the best possible estimates.

##### Effect of co-solutes

To evaluate the effect on RNA phase separation, co-solutes were added to 3.75 g/L RNA with 10 mM $MgCl_2$ in a 96-well plate at the concentrations indicated. The samples were mixed by shaking by hand and the degree of phase separation was quantified by measuring optical density at 500 nm using a Tecan Pro plate reader.

##### Confocal Microscopy

###### Partitioning of small organic molecules

The fluorescence images related to the partitioning of small molecules were acquired and analyzed using a Spinning Disk Confocal (Nikon, Andor CSU) microscope with ×40 or ×63 oil immersion lens (University of Perugia). Fluorescence images related to the partitioning of short chain fatty acids, phospholipids and populations experiments were acquired by using and analyzed using a FluoView FV1200 (Olympus) confocal laser scanning microscope (University of Okayama). Confocal images were acquired for more than 5 independent experiments with similar results. The images collected were analyzed using ImageJ.

###### Partitioning of fatty acids and phospholipid like molecules

RNA condensates were prepared as previously described using a saturated RNA solution. Phospholipid ($\beta$-BODIPY C12-HPC, Hexadecanoyl-sn-Glycero-3-Phosphocoline) or short chain fatty acids (we tried both C11 undecylenic acid and C12 dodecanoic acid) were added 5 - 10 µM after RNA condensate formation with 10 g/L RNA in the presence of 100 mM

$MgCl_2$ or HCl at 25 °C. After few minutes, around 30 uL were placed on a glass slide along with a cover slip. The experiments last for 2 - 4 hours.

**pH measurements within RNA droplets**

LysoSensor Yellow/Blue DND-160 (Thermo Fisher Scientific, catalog number L7545) was used as a pH-sensitive fluorescent probe for pH measurements within RNA condensates. Working solutions were prepared at a final concentration of 20 µM at 25 °C. Calibration measurements were performed across pH 1.67 - 5.50 using Tris-HCl buffer at varying concentrations (0 mM, 10 mM, 25 mM, 50 mM, 75 mM, 100 mM). For each pH point, LysoSensor (20 µM final concentration) was added and fluorescence intensity was measured by fluorescence microscopy using 400 nm excitation. The calibration curve was generated by plotting $\log_{10}$ (fluorescence intensity at 400 nm) versus measured pH. RNA condensates were formed using yeast RNA (7.5 g/L) and $MgCl_2$ (50 mM) in solutions buffered to different pH values using Tris-HCl buffer. LysoSensor probe (20 µM final concentration) was added after condensate formation. Fluorescence microscopy was performed using an Andor spinning disk microscope with 400 nm excitation. For quantitative pH measurements, regions of interest (ROIs) were manually defined around individual condensates, and mean fluorescence intensity was measured for each ROI. Multiple droplets were analyzed for each external pH condition. Internal pH values were calculated by applying the calibration curve equation to the measured fluorescence intensities. Data are presented as individual droplet measurements to illustrate the systematic pH difference between condensate interior and external solution.

**FRAP Experiments**

The fluorescence recovery after photobleaching (FRAP) experiments were carried out using a laser confocal system Nikon A1R System on a Nikon TiE2 inverted microscope using an 100x oil objective (100×/1.49 Oil Apo TIRF). Rhodamine 6G (R6G, 35 µM) was used as a fluorescent tracer for FRAP analysis. RNA condensates were formed using total yeast RNA (7.5 g/L) and varying the $MgCl_2$ and HCl concentrations (both 10 mM final concentration).

The 561 nm laser line was used for both fluorescence photobleaching and detection. The fluorescence decay and recovery were measured using a GaAsP detector (Nikon DUB4). All the experiments were designed and carried out using Nikon NIS Elements software. The data were exported as single Excel file to be then processed in a MATLAB-based routine. The recovery time (seconds) was extracted from the fitting of normalized time-dependent intensity curves using a least squares regression. Imaging array and pixel resolution were optimized to obtain a sampling ratio between 4 and 15 frames per second depending on how fast the kinetics were considered in this study. An average bleaching spot of 1.5 microns in diameter was obtained throughout the experiments with a bleaching duration ranging between 115 and 215 msec. The temperature was kept constant at 25 °C. The imaging settings were adjusted such that at least 500 - 1000 images could be acquired without an overall bleaching of more than 10% of the initial signal. The bleaching experiments consisted of three phases starting with the acquisition of 20 - 100 pre-bleach images to determine the steady state pre-bleach value of the fluorescence signal and to ensure that the focal plane was maintained during the imaging. A reasonably short bleach spot was then applied approximately in the middle of the target droplet. The depth and precision of the bleached region were determined by a simple line measurement across the bleached region. The bleach duration was minimized but a bleaching efficiency of no less than 85% was maintained. Typically, 10 - 15 droplets from the same batch were imaged for each experiment (total duration <50 min to prevent artifacts due to the aging of the droplets). The droplets were freshly prepared before each set of measurements. The volume containing droplets were seeded in a small chamber (200 µL) glued on a coated fluorescence microscopy suitable coverslip. After a short settling time the kinetic measurements were initiated (generally 2 - 4 minutes).

**RNA condensate populations experiments**

RNA condensates were prepared from a saturated RNA solution (7 - 10 g/L) by adding 100 mM HCl or 100 mM $MgCl_2$. Before triggering phase separation, fluorescently labeled (FAM, Cy3 and Cy5) oligos were added to the RNA solution at the final concentration of 750 nM. In detail, the following 20 nt sequences for ssRNA (FAM or Cy5 labeled) or ssDNA (FAM or Cy3 labeled) were used. dsDNA was prepared by annealing a FAM-labeled oligo and its unmodified complement. They were ordered as separate oligos from IDT and annealed at 10 µM in 50 mM Tris-HCl (pH 8) with 100 mM NaCl (98 °C, 2 minutes and ramping to 37 °C at 0.1 °C per second). Oligo sequences are given in Table 1. Different populations (with or without the addition of a single fluorescently labeled ssRNA or ssDNA) were prepared in separate glass tubes (10 mL final volume). After 2 - 3 hours of incubation in the dark at 25 °C the condensates were gently resuspended by pipetting and mixed together in another microcentrifuge tube. After 5 minutes, around 30 µL of mixed solution containing two populations was placed on the glass slide along with a cover slip for visual analysis.

Table 1: Oligonucleotides used for fluorescence microscopy.

| **Name** | | **Sequence 5' to 3'** |
|---|---|---|
| ssRNA | | /56-FAM/AAGCUUGGUCAGGCUUGUAC |
| | or | /5Cy5/AAGCUUGGUCAGGCUUGUAC |
| | or | /56-FAM/aaaaaaaaaaaaaGGCAUCCUGGAUUCCACUCGCC |
| ssDNA | | /56-FAM/AAGCTTGGTCAGGCTTGTAC |
| | or | /5Cy3/AAGCTTGGTCAGGCTTGTAC |
| dsDNA | | /56-FAM/AGGAGAGGAGAGAGAAGAGATCCT |
| | and | AGGATCTCTTCTCTCTCCTCTCCT |

#### Small organic molecule catalysis within RNA condensates

##### Addition reaction

Fresh stock solutions of 2,4-DNPH (116 mM) and aldehydes (300 mM) were prepared in water under acidic condition. Addition reaction between 2,4-DNPH (200 µM) and the aldehydes (p-anisaldehyde and vanillin, final concentration 400 µM) were performed by preparing the condensates using 5 g/L RNA, at 25 °C, using 100 mM $MgCl_2$ to trigger phase separation. Please note that the addition reaction using RNA condensates prepared with strong acids such as HCl leads to a too fast reaction under this experimental condition. The formation of C-N bonds was monitored by recording the absorbance spectra from 240 to 800 nm in a final volume of 200 µL using a MultiSkan SkyHigh microplate reader with a 96-well UV-transparent plate. The reaction was monitored under three different conditions: in water, condensate, and supernatant phase. The experiments using the supernatant phase were performed by preparing the condensates as reported above in this section. The solution was then centrifuged for 30 minutes at 16000 g for 30 minutes. The supernatant phase has been separated from the condensate phase and used for the experiments. Since 2,4-DNPH has been reported to interact with protein and nucleic acids, we checked the stability of the 2,4-DNPH signal over time (total time of 60 minutes) in the presence of RNA. Within 60 minutes, the absence of non-specific DNPH signal between 2,4-DNPH and RNA was recorded (although after a few hours it is possible to measure a change in the maximum and intensity of the absorbance peak, indicating interactions between RNA and the hydrazinic compound; Fig. S12c). At this point, we set up the experiment as follows: first, we added RNA 5 g/L, then 2,4-DNPHZ (200 µM) and immediately after aldehydes (400 µM) with $MgCl_2$ 100 mM at 25 °C. In the water samples, pH was adjusted using HCl to match the pH of the RNA condensate and supernatant. The spectra were recorded every 2 - 5 minutes and the signal reported in the kinetic analysis is the decrease in the absorbance of the peak related to 2,4-DNPHZ.

**pNPA hydrolysis**

Fresh stock solution of pNPA (200 mM) were prepared in DMSO 100% and pNPA hydrolysis was evaluated at three different concentrations (4 mM) and (1 mM) in 1 mL of total solution at 25 °C in water, condensates, and supernatant. Acid hydrolysis of pNPA is slower than that at neutral or basic pH. In addition, the solubility of pNPA is limited. For this reason, we used a total RNA concentration equal to 5 g/L to prepare the condensate phase and the supernatant in the presence of 100 mM $MgCl_2$ and conducted the experiments for longer times (around 360 minutes). The same experiment was performed using condensates prepared by adding different HCl concentrations (decreasing the pH further by adding acid increases the reaction rate related to pNPA hydrolysis). The detection of pNP under acidic conditions is through the monitoring of a band around 320 nm which is completely covered by the RNA signal in our samples and has a weak absorbance signal (protonated form). For this reason, we decided to monitor the signal of the formed pNP at pH 8.0 which is completely deprotonated. As reported in Fig. S16 we have used KP 200 mM pH 8.0 as buffer to convert pNP in the deprotonated form. Briefly, 50 µL of the reaction mix under acidic conditions have been placed on a 96-well plate along with 150 µL of KP 200 mM pH 8.0 at each time point and immediately read at 405 nm using the MultiSkan SkyHigh microplate reader.

**Glycine formation**

The RNA condensates have been prepared by using 100 mM $MgCl_2$ in the presence of 7.5 g/L RNA. All components of the reaction mixture have been added before $MgCl_2$ which triggers the condensation of RNA. In detail, we used a final concentration of 20 mM for glyoxylate, and 10 mM for l-amino acids with a multichannel pipette to start the reaction at the same time in a final volume of 1 mL under agitation (360 minutes total experimental time). The reaction has been performed at 25 °C and at different time points we have withdrawn 2.5-10 µL and placed in 50 µL borate buffer 0.2 M with 100 µM L-norvaline and then derivatized as described in the supporting information **"Derivatization of glycine and other amino acids coupled with HPLC analysis"**. The experiments using the supernatant phase were performed by preparing the condensates as reported above in this section. The RNA condensate solution was then centrifuged for 30 minutes at 16000 g for 30 minutes. The supernatant phase has been separated from the condensate phase and used for the experiments.

**Alanine formation** The alanine formation experiments from pyruvate had to be performed under milder conditions compared to glycine formation. Higher concentrations of pyruvate (higher than 10 mM) combined with glycine/diglycine (higher than 8 - 10 mM) in the presence of 20 - 40 mM $MgCl_2$ prevented formation of liquid droplets and instead produced solid aggregates (the formation of the aggregates was time dependent. Therefore, conditions were optimized to maintain RNA condensate properties while enabling detectable transamination. The optimized conditions were: 25 mM $MgCl_2$ in the presence of 5 - 7.5 g/L RNA, with 9.5 mM pyruvate as the $\alpha$-ketoacid, and 7.5 mM for glycine and diglycine (or 1 mM for hexaglycine). All components of the reaction were mixed before addition of $MgCl_2$ which triggers the condensation of RNA. The transamination reactions were initiated simultaneously using a multichannel pipette in a final volume of 1 mL under agitation (360 minutes total experimental time). The reaction has been performed at 25°C and at different time points 5 - 10 µL samples were withdrawn and placed in 50 µL borate buffer 0.2 M with 100 µM L-norvaline. Derivatized was performed as described in the supporting information under **"Derivatization of glycine and other amino acids coupled with HPLC analysis"**. Despite these milder reaction conditions, necessitated by the requirement to maintain proper RNA phase separation, significant enhancement of glycine-to-alanine transamination was observed in RNA condensates, with approximately 7-fold enhancement in initial reaction rates compared to water controls and 4 - 6-fold higher final alanine concentrations. The experiments using the supernatant phase were performed by preparing the condensates as reported above in this section. The RNA condensate solution was then centrifuged at 16000 g for 30 minutes. The supernatant phase has been separated from the condensate phase and used for the experiments. It should be noted that overall alanine production remains relatively low under these conditions due to the milder reaction conditions, the inherently lower reactivity of pyruvate compared to glyoxylate, and the reduced number of reactive droplets formed under the optimized conditions.

##### Enzymatic reaction within RNA condensates

##### Cystalysin activity assay

Cystalysin from *Treponema denticola* is a pyridoxal 5-phosphate (PLP)-dependent enzyme that catalyses the $\alpha,\beta$-elimination of $\beta$-chloroalanine to pyruvate, ammonia and HCl. Since this enzyme produces HCl, it can trigger phase separation of RNA by lowering the pH of the entire water solution. Cystalysin has been expressed and purify as previously described.[2] The activity of cystalysin in the presence of RNA was checked by monitoring the decreasing in pH and the pyruvate formation by using LDH coupled assay as previously reported for this enzyme[3] and for others.[4] Briefly, before to start the experiments showed in the Fig. S19, we have verified the stability and the activity of the protein in water at pH 5.5 in the presence of RNA and $MgCl_2$ at different concentrations (from 5 mM to 100 mM at 25 °C) (data not shown). Pyruvate formation during the reaction of cystalysin with $\beta$-chloroalanine was assayed by coupling the pyruvate produced to the NADH-dependent lactate dehydrogenase-catalyzed reaction. The assays were performed at 25 °C and the NADH signal has been measured by using a MultiSkan SkyHigh microplate reader and UV-transparent 96-well plate. A typical reaction mixture contained 1 mM $\beta$-Cl-alanine, 300 µM NADH, 1-5 µM lactic acid dehydrogenase in water in the presence of RNA 400 ng/µL, in a final volume of 200 µL. The reaction was initiated by addition of several concentrations of cystalysin 0.5 - 2 µM. A molar extinction coefficient of $\varepsilon_{340} = 6220\,\mathrm{M^{-1}cm^{-1}}$ for NADH was used. Determination of the kinetic parameters for the reactions catalyzed by cystalysin was performed by measuring the initial velocities by the assay described above.

**OxDC activity assay** OxDC wild type has been expressed and purified as previously reported.[5,6] The enzymatic activity of OxDC within the condensates was measured by detecting the product formate using potassium permanganate as a chemiluminescent reagent. This approach has been extensively validated for formate detection in our previous work.[5] **Assay optimization and detection method**. The detection system was initially calibrated using different formate concentrations at a fixed potassium permanganate concentration of 1 mM. Optimal sensitivity was achieved by monitoring the absorbance variation at 545 nm, consistent with our previously established protocol.[5] This wavelength corresponds to the characteristic absorption of the permanganate-formate reaction product, providing a reliable and quantitative measure of formate concentration. **Condensate preparation and reaction conditions**. RNA condensates were prepared in Tris-HCl 32 mM pH 7.5 buffer by mixing 3.7 g/L of RNA with OxDC at concentrations of 0.2 µM and 0.4 µM in the presence of $MgCl_2$ 50 mM. The substrate oxalate was added to a final concentration of 10 mM, and all reactions were performed at 25°C under controlled conditions. **Reaction termination and detection protocol**. The enzymatic reaction was terminated by rapidly mixing 25 µL of the reaction mixture with 175 µL of potassium phosphate buffer 100 mM pH 8.0 containing 1.75 mM potassium permanganate in a total volume of 200 µL. The absorbance at 545 nm was immediately monitored for the first 5 - 10 minutes using a MultiSkan SkyHigh microplate reader maintained at 25°C. This timing is critical as it captures the initial linear phase of the permanganate-formate reaction before potential interference from secondary reactions. **Data analysis**. All measurements were performed at least in triplicate, and the data were analyzed using GraphPad Prism 10.4.1 (GraphPad Software, Boston, Massachusetts USA, www.graphpad.com). Formate concentrations were determined using standard curves generated with known formate concentrations under identical buffer conditions.

##### Ribozyme activity

We used a hammerhead ribozyme that had been shown to be active inside coacervate samples.[7] The sequences used are given in Table 2. The substrate, including a polyA spacer, was ordered as solid-phase synthesized RNA from FASMAC. DNA template encoding the hammerhead ribozyme was prepared by annealing synthetic DNA oligos (Eurofins) **'Ribozyme DNA template forward'** and **'Ribozyme DNA template reverse'** to each-other (98 °C for 2 minutes and ramping to 37 °C at 0.1 °C per second using 10 µM oligo each, 100 mM $MgCl_2$, and 100 mM NaCl). RNA was generated using ScriptMAX Thermo T7 Transcription Kit (TOYOBO, TSK-101) according to the manufacturers instructions using 1 µL of the annealed oligos as the template. After a DNA digest (DNase I, Takara, 2270A), the ribozyme RNA was silica column-purified using the RNA Clean and Concentrator-5 (Zymo,

R1014) and eluted in RNA-grade water. The concentration and purity were determined by Nanodrop. Reactions were set up in using 50 mM Tris-HCl (pH 3 in the absence of yeast RNA or pH 7 in the presence of yeast RNA), 2.5 mM KCl, 0.2 µM substrate as well as $MgCl_2$, yeast RNA, and ribozyme in amounts indicated. Reactions took place at 37 °C for 30 minutes. The reactions were either precipitated with ethanol and sodium acetate (in cases without phase separation) or separated into supernatant and condensate phase by centrifugation at 18 000 g for 5 minutes. Both the condensate phase and precipitated reactions were resuspended in TBE-Urea Sample Buffer / RNA loading dye (Novex) at 95 °C for 2 minutes, cooled on ice, and completely loaded onto a 15% TBE-polyacrylamide gel containing 8 M urea. Ribozyme activity was estimated after PAGE by quantifying the cleaved bands relative to the uncleaved substrate band on pictures taken with the Typhoon FLA 7000 (GE) using ImageJ.

Table 2: Sequences used for the study of ribozyme activity. The T7 promoter (including the first three Gs) is underlined.

| **Name** | **Sequence 5' to 3'** |
|---|---|
| Ribozyme substrate | /56-FAM/aaaaaaaaaaaaaGGCAUCCUGGAUUCCA-CUCGCC |
| Ribozyme sequence | GGGCGAGGUACAUCCAGCUGACGAGUCCCAA-AUAGGACGAAAUGCC |
| Ribozyme DNA template forward | <u>TAATACGACTCACTATAGGG</u>CGAGGTACATCC-AGCTGACGAGTCCCAAATAGGACGAAATGCC |
| Ribozyme DNA template reverse | GGCATTTCGTCCTATTTGGGACTCGTCAGCTG-GATGTACCTCG<u>CCCTATAGTGAGTCGTATTA</u> |

#### Statistics and reproducibility

The experimental data reported in this manuscript were performed at least in triplicate. The error bars shown along the manuscript represent the SD. Data were analyzed with GraphPad

Prism 10.4.1 (GraphPad Software, Boston, Massachusetts USA, www.graphpad.com). The authors will provide all the necessary data to evaluate the manuscript.

#### Characterization and identity of phase-separated RNA

We were interested in what species of RNA the commercially available yeast RNA preparations contain. However, the vendors (Roche and Wako) were unable to provide details upon request, citing confidentiality in the manufacturing process. To determine at least the size distribution of the RNA in the preparations, denaturing PAGE was performed. The results show that both RNA preparations contained mostly small RNAs <50 nt in size with the majority being smaller than 20 nt (Fig. S1b).

After inducing phase separation by adding 100 mM $MgCl_2$, RNA condensates were separated from the supernatant by centrifugation and PAGE was performed on the obtained droplet and supernatant fractions. The mobility inside the polyacrylamide gel indicates that the pellet is enriched in longer (up to around 50 nt) RNA fragments while the supernatant mostly contains short (<20 nt) RNA fragments (Fig. S1b). We also noticed that the RNA phase is very viscous and hard to resuspend. Microscopic observations support the assumption of a high viscosity because compared to other LLPS systems the RNA condensates do not readily coalesce (for example Fig. S6c). Additionally, the RNA size distribution of RNA preparations from different vendors before and after phase separation are similar (Fig. S1b). We assume that RNA is composed of fragments produced from yeast RNA through random hydrolysis with a more or less random sequence. However, the exact sequence composition is unknown.

Atomic emission spectroscopy was performed to characterize elements present in the RNA preparations. Only phosphorous (from the RNA backbone) and traces of magnesium (around 200 µM were found in 1 mL of the RNA stock solutions (5 g/L), see supplementary table available as a separate excel file).

#### The use of commercial yeast RNA as a surrogate

One important issue of our system is the unknown sequence and length composition of the yeast RNA used. Yeast RNA contains mature, processed transcripts with modifications and specific evolutionary-derived sequences. It probably differs substantially from the prebiotic scenarios. This creates uncertainty about whether similar phase separation behavior would occur with truly prebiotic RNA populations. Without knowing which RNA sequences drive the phase separation behavior, it is difficult to establish clear molecular mechanisms and structure-function relationships. This limitation makes it impossible to determine the precise conditions under which RNA condensation would occur prebiotically and which specific RNA features (length, secondary structure, sequence motifs) are critical for the observed behaviors. However, the competing characteristics of the backbone (more hydrophilic) and nucleobases (more hydrophobic) are a conserved feature of all nucleic acids. We therefore assume that the phase separation process is sufficiently robust to occur under many different scenarios of RNA length, sequence composition, and environmental parameters.

#### Hydrolysis of pNPA under acidic conditions

Acid-catalyzed hydrolysis of pNPA (para-nitrophenyl acetate) is significantly less favorable than base hydrolysis due to several mechanistic factors. Acid hydrolysis requires protonation of the ester carbonyl followed by nucleophilic attack by water, which is inherently a weak nucleophile, particularly in acidic environments where hydroxide ion concentration is minimal. The reaction must proceed through a tetrahedral intermediate that requires stabilization. Moreover, the strongly electron-withdrawing para-nitro group hinders carbonyl protonation a critical step in acid hydrolysis—under acidic conditions. The nucleophilic attack by water faces a substantially higher energy barrier compared to hydroxide attack, and although carbonyl protonation helps reduce this barrier, water remains a weak nucleophile. These mechanistic limitations manifest in kinetic constants that are orders of magnitude lower for acid hydrolysis compared to base hydrolysis. In acidic environments (pH 2-3), the rate constant k typically ranges from $10^{-6}$ - $10^{-7}$ $s^{-1}$, while in basic environments (pH 8-9), k values are approximately $10^{-3}$ - $10^{-4}$ $s^{-1}$. This 3 - 4 orders of magnitude difference explains our experimental observations at pH 2.0 - 2.5, where conversion reached only 1 - 1.5% after 6 hours. It also contextualizes why the catalytic effect of RNA condensates, translates to a relatively modest acceleration in absolute terms.

To comprehensively evaluate the catalytic effect of RNA condensates on pNPA hydrolysis, we conducted experiments using 1 and 4 mM pNPA under three distinct conditions. Notably, pNPA at 4 mM concentration typically exhibits solubility limitations and tends to crystallize. However, the combination of high RNA content and 5% DMSO in our experimental setup enhanced pNPA solubility through multiple mechanisms: hydrophobic interactions, preferential partitioning within RNA condensates (which possess physicochemical properties distinct from the aqueous environment), and hydrogen bonding. These solubilizing effects could also contribute to our observed enhanced hydrolysis rates in RNA condensates. Following rigorous model comparison, we determined that a zero-order kinetic model provided a better fit for our experimental data ($R^2 > 0.96$) across all conditions) compared to first-order kinetics. This aligns with the extremely low conversion rates ($\leq 1.5\%$ after 6 hours), which create conditions where substrate concentration remains effectively constant throughout the reaction period. Additionally, at pH 2.5, the rate-limiting step may operate independently of substrate concentration.

For precise quantification, we converted absorbance measurements to concentration using $\varepsilon_{365} = 18\,000\,\mathrm{M^{-1}cm^{-1}}$ for deprotonated pNP at pH 8.0 (calculated based on the microplate pathlength with a total volume of $200\,\mu\mathrm{L}$). To ensure measurements within the linear range of detection, samples were diluted 2-fold and 4-fold for 1 mM and 4 mM experiments, respectively, prior to analysis. Linear regression analysis of concentration versus time data yielded slopes (k values) and $R^2$ values for each experimental condition, enabling calculation of relative rates and concentration-normalized efficiencies. As detailed in Table of Fig. S16, RNA condensates demonstrated a modest but consistent catalytic effect, corresponding to a 1.09-

fold acceleration versus water at 1 mM pNPA and 1.14-fold acceleration at 4 mM pNPA. Statistical analysis incorporated mean values from experimental replicates with standard deviations to quantify experimental variability.

#### Addition reaction under acidic conditions

This study investigated the kinetic behavior of hydrazone formation reactions between 2,4-dinitrophenylhydrazine (2,4-DNPH) and two different aromatic aldehydes (p-anisaldehyde and vanillin) in three distinct reaction environments: RNA condensates, diluted phase (containing RNA and $Mg^{2+}$), and acidic water (pH 2.0 - 2.5). The primary questions addressed were:

- How do RNA condensates affect the kinetics of hydrazone formation compared to the diluted phase?
- How do the reaction rates in both environments compare to standard aqueous conditions?
- What is the mechanism responsible for any observed rate enhancements in RNA condensates?

All reactions were carried out around pH 2.0 - 2.5, with initial concentrations of 200 µM for 2,4-DNPH and 400 µM for aldehydes. A second-order kinetic model was applied to determine the rate constants and half-lives. The reaction follows second-order kinetics for unequal initial concentrations ($[A]_0 \neq [B]_0$). This model provided an excellent fit to the experimental data ($R^2 > 0.95$ for all conditions), validating the second-order kinetic behavior.

The rate constants observed in acidic water (189 $M^{-1}min^{-1}$ for p-anisaldehyde and 104 $M^{-1}min^{-1}$ for vanillin) align well with literature values for similar reactions at comparable pH, which typically range from 20-200 $M^{-1}min^{-1}$.[8,9] The observed ratio of reactivity between p-anisaldehyde and vanillin (1.81-fold in acidic water) is consistent with established structure-activity relationships, which predict higher reactivity for para-methoxy substituted aldehydes compared to those with meta-methoxy groups.[9–11]

The close correspondence between the concentration enhancement factor (6.0) and the reaction acceleration factors (6.7-8.8) suggests that the mechanism for rate enhancement in RNA condensates is a local increase in the concentration of reactants. This provides a clear physicochemical explanation for the catalytic effect of RNA condensates and is consistent with recent literature on reaction acceleration in phase-separated environments.[12,13] Several mechanistic features support this conclusion:

- Concentration effect: the measured 6-fold higher concentration of 2,4-DNPH in RNA condensates closely matches the observed rate enhancements.
- Preservation of reactivity ratios: the relative reactivity of p-anisaldehyde versus vanillin remains consistent across environments (1.8 - 2.6), suggesting the intrinsic reaction mechanism remains unchanged.
- Kinetic model consistency: the second-order kinetic model fits well across all environments, indicating that the same reaction mechanism operates in both condensates and dilute phase.

The slightly higher enhancement factor for p-anisaldehyde (9-fold) compared to vanillin (7-fold) could also reflect subtle differences in partitioning behavior between the two aldehydes. This could be attributed to vanillin's additional hydroxyl group, which increases its hydrophilicity (logP $\approx$ 1.2 vs. 1.8 for p-anisaldehyde)[14] and may affect its distribution between the RNA-rich and RNA-poor phases.[15] Also, the hydroxyl group of vanillin likely forms hydrogen bonds with solvent molecules or reaction intermediates, altering approach trajectories and stabilization of transition states.[8,16] Overall, these findings have some important implications in prebiotic chemistry. Indeed, RNA condensates could have served as primitive compartments that accelerated bimolecular condensation reactions relevant to early chemical evolution[17] by increasing the local concentration of small molecules.[18,19]

#### Transamination under acidic conditions: mechanistic implications

Non-enzymatic transamination between an amino acid and an $\alpha$-keto acid (like glyoxylate, CHO-COOH) typically proceeds through the following steps. Initial Schiff base formation in which the nucleophilic amino group of the amino acid attacks the electrophilic carbonyl carbon of the $\alpha$-keto acid (generating a carbinolamine intermediate — hemiaminal). Then, dehydration occurs eliminating water to form the imine (Schiff base). At this point, the imine undergoes tautomerization (proton shift) and a proton transfers from the $\alpha$-carbon of the original amino acid to the nitrogen-carbon double bond (ketimine intermediate). Finally, the ketimine undergoes hydrolysis through nucleophilic attack by water which generates a new $\alpha$-keto acid (from the original amino acid) and a new amino acid (glycine in the case of glyoxylate).

Under conditions of acidic pH, the amine groups of the amino acids are protonated, reducing their nucleophilicity and rendering transamination highly inefficient (rate limiting step). Optimal pH (transamination is highly pH dependent) is typically near neutral (6 - 8) where amino groups retain some nucleophilicity while the carbonyl is sufficiently electrophilic. Importantly, at low pH glyoxylate is predominantly in its hydrated form. The hydrated form (geminal diol) of glyoxylate inhibits transamination because it lacks the electrophilic carbonyl group necessary for nucleophilic attack by amines. This is particularly problematic under acidic conditions, where the equilibrium strongly favors the hydrated form, requiring an energetically unfavorable dehydration step before transamination can proceed.

Besides pH, there are other challenges related to non-enzymatic transaminations. One challenge is the lack of precise orientation: without enzyme active sites orienting the reactants, reactants meet in random orientations and this reduces the probability of productive collisions. In addition, competing/side reactions such as deamination of amino acids (which is occurring, data not shown), racemization, aldol condensations of the $\alpha$-keto acid, and polymerization reactions could occur, limiting the reactant concentration.

RNA condensates could enhance non-enzymatic transamination through several mechanisms. For example, RNA condensation creates a microenvironment with different properties compared to the bulk solution (pH, water activity, etc. that might favor amine deprotonation) or transition state stabilization where the RNA structure may stabilize charged intermediates. The activation barriers, particularly for the rate-limiting steps, could thus be lowered by RNA (RNA functional groups can serve as proton donors and acceptors). Highly concentrated RNA may also provide sterically confined regions that position reactants in favorable orientations, thereby reducing the entropic penalty for bringing reactants together. Further, RNA binds metal ions, like $Mg^{2+}$, which could create catalytic centers by coordinating reactants and facilitating electron transfers (enhancing glyoxylate electrophilicity through coordination, stabilizing RNA structures, forming coordination complexes with amino acids, particularly l-ornithine and l-lysine, and providing Lewis acid catalysis even in the dilute phase).

In summary, non-enzymatic transamination proceeds through classic organic chemistry mechanisms but faces significant challenges in terms of efficiency and selectivity compared to enzymatic processes, especially under acidic conditions. The remarkable enhancement observed in the presence of RNA condensates (Main Fig. 5c and Fig. S13) suggests that even in the absence of enzymatic catalysts adapted by evolution, complex macromolecular environments can substantially facilitate this fundamental chemical transformation. These results can prove insight into how metabolic reactions might have occurred in early prebiotic scenarios. Additionally, the apparent role of magnesium indicates how metal ions could have been essential co-catalysts in prebiotic systems.

#### Transamination under acidic conditions: kinetic data

We have calculated reaction rates by applying linear regression to the first three time points (0, 10, and 30 minutes) of glycine formation for each amino acid and condition. The slope of this regression line represents the initial reaction rate. The initial reaction rates show remarkable enhancement in condensates:

- Lysine: 19-fold enhancement over water

  38 µM/min (condensates) vs 7.0 µM/min(supernatant) vs 2 µM/min (water)

- Alanine: 14-fold enhancement

  14 µM/min (condensates) vs 4 µM/min (supernatant) vs 1 µM/min (water)

- Ornithine: 11-fold enhancement

  107 µM/min (condensates) vs 14 µM/min (supernatant) vs 10 µM/min (water)

The maximum glycine yields follow the same pattern:

- Lysine - 16.7-fold relative enhancement (condensates vs water)
- Alanine - 8.6-fold enhancement
- Ornithine - 8.7-fold enhancement

All amino acids show efficiency ratios of 3.5 - 7.6-folds (condensates vs supernatant). This exceeds what can be explained by concentration differences alone, because amino acids were not enriched in the condensates (Fig. S10). RNA alone shows an effect, because supernatant (containing dissolved short RNA) shows 1.5 - 4.5-fold higher rates than water. Condensates provide additional 3 - 8-fold enhancement over supernatant. The data are reported in Fig. S20.

#### Transamination reaction under acidic conditions: products of transamination

When glyoxylate (CHO-COOH) undergoes non-enzymatic transamination with amino acids like l-alanine, l-lysine, and l-ornithine in an RNA-$Mg^{2+}$ droplet system under acidic conditions, the following products can form depending on which amino group participates in the reaction. $\alpha$-amino group transamination products:

- glyoxylate + l-alanine → glycine + pyruvate
- glyoxylate + l-lysine → glycine + 2-keto-6-aminocaproic acid
- glyoxylate + l-ornithine → glycine + 2-keto-5-aminovaleric acid

or terminal amino group transamination products (specific to lysine and ornithine):

- glyoxylate + $\epsilon$-$NH_2$ group of l-lysine → glycine + modified lysine with altered $\epsilon$-$NH_2$ group (5-aminopentanal)
- glyoxylate + $\delta$-$NH_2$ group of l-ornithine → glycine + modified ornithine with altered $\delta$-$NH_2$ group (4-aminobutanal)

In the RNA and $Mg^{2+}$ condensate system, the distribution of these products may differ from what would be expected in solution due to several reasons: the catalytic effects of $Mg^{2+}$ as a Lewis acid, electrostatic interactions between RNA phosphates and protonated amino groups, altered local concentration of reagents within the condensates, or potential RNA-mediated positioning of reactants. The products retain their side chain amino groups, allowing detection via DNFB derivatization (however, they can undergo cyclization). Indeed, other peaks were seen during HPLC determination of glycine in presence of l-ornithine and l-lysine.

#### Qualitative determination of pNPA, pNP and 2,4 DNPH through HPLC

pNPA, pNP and 2,4-DNPH were also determined using HPLC (Fig.S16). 20 µL of the reaction mix were mixed with 20 µL 60/40% ACN/water (v/v) and the mixture was centrifuged for 10 minutes to remove the precipitated RNA. Then, 20 µL were loaded on an EC NUCLEODUR 100-5 C18 (250 × 4.6 mm) Machery-Nagel column at 25 °C connected to a JASCO LC-4000 HPLC control system. The mobile phase was 60/40% ACN/water (v/v) the run was performed at a flow rate of 1 mL/min. A Jasco UV-4070 detector set at 360 nm was employed. The peaks corresponding to the different compounds were analyzed using the ChromNAV software (version 2.4.0.5). Standards have been used to confirm the retention times of the compounds.

#### Derivatization and HPLC analysis for glycine and other amino acids

Amino acid concentrations were measured by HPLC after derivatization with 1-Fluoro-2,4-dinitrobenzene (DNFB) according to.[20,21] Briefly, 5-10 µL of reaction mix was placed in 40-45 µL of borate buffer 0.2 M pH 9.0. As an internal control, 100 µM L-Norvaline was added to each sample in a final volume of 50 µL. After the addition of 25 µL of DNFB 72 mM to each mixture, the samples were incubated at 60 °C for 1 h in the dark. The reaction was then stopped on ice by adding 175 µL of cold PBS pH 7.0 (final volume 250 µL). The samples were then centrifuges at 18000 g for 20 minutes and stored at -20 °C before HPLC analysis. 20 µL of derivatized samples were loaded onto a EC NUCLEODUR 100-5 C18 (250 × 4.6 mm) Machery-Nagel column at 25 °C connected to a JASCO LC-4000 HPLC control system. The mobile phases were as follows: A, 40 mM $NaH_2PO_4$ buffer pH 7.8; B, 45/45/10% ACN/MetOH/water. The conditions used for the elution are the same as reported[22] and the run was performed at a flow rate of 1 mL/min. A Jasco UV-4070 detector set at 360 nm was employed. The peak corresponding to the different amino acids was integrated using ChromNAV software (version 2.4.0.5). A standard curve for peak area estimation was prepared using the commercially available amino acids used in the study.

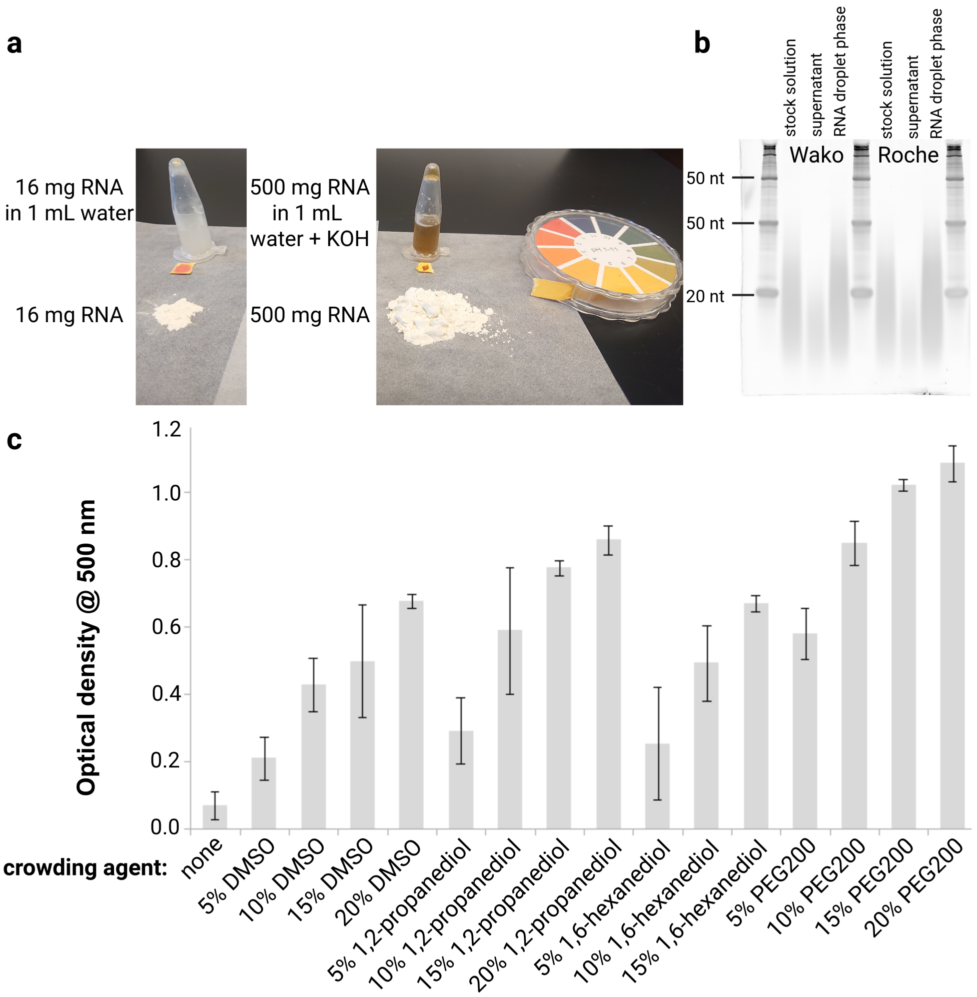


Supplementary Figure S 1: **RNA solubility, size distribution, and effect of crowding agents on phase separation**. **a,** RNA powder is somewhat soluble in pure water, but the solution of 16 mg in 1 mL water is milky and acidic, indicating incomplete dissolution (left). Addition of a base makes the solution transparent again (not shown). Upon repeated addition of KOH to the RNA solution, it was possible to dissolve 500 mg of RNA in a total volume of 1 mL with the solution becoming progressively more viscous and brownish but staying acidic (right). The pH of the RNA solution stayed acidic after the addition of KOH as confirmed by spotting the solution of pH paper. **b,** PAGE gel showing the size distribution of the RNA used. The size marker is DynaMarker RNA Low II (BioDynamics Laboratory). RNA from two different vendors had the same composition of mainly short polynucleotides of <50 nt in length. After triggering phase separation by addition of 100 mM $MgCl_2$ (to 4 g/L RNA final) and separating the droplet and supernatant by centrifugation, the supernatant contained mainly short RNAs (<20nt) while the droplet phase contained the longer RNA species. **c,** Effect of crowding agents on phase separation of 2.5 g/L RNA in the presence of 10 mM $MgCl_2$ and 3.3 mM bis-tris-propane (pH 7.2). The addition of DMSO, 1,2-propanediol, 1,6-hexanediol, and PEG200 increased the phase separation in a dose-dependent manner. In the presence of 20% PEG200 phase separation occurred even without any added magnesium (not shown, but similar to reported data).[23]

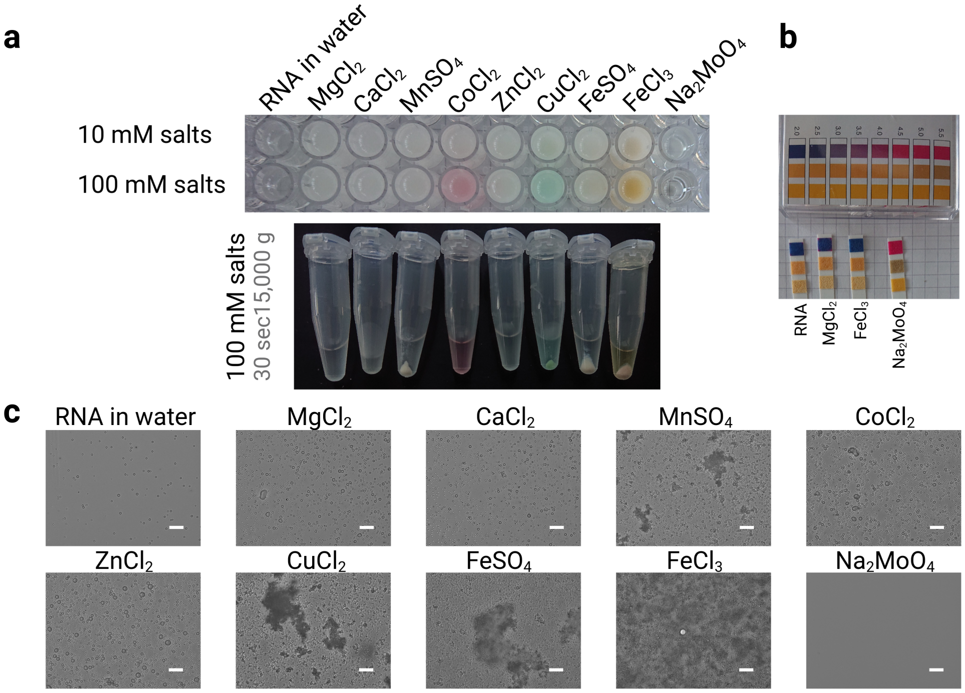


Supplementary Figure S 2: **Observation of RNA phase separation in the presence of different multivalent salts**. **a,** The RNA solution was supplemented with 10 mM (top row) or 100 mM (bottom row) of different salts (dissolved as 1 M stocks in water). The solution turned whitish indicating phase separation in all cases except for sodium molybdate. The samples prepared with 100 mM salts were spun down to visualize the pellets formed. **b,** pH paper indicating that the pH of the sodium molybdate-RNA mixture increased compared to pure RNA, RNA-$MgCl_2$, and RNA-$FeCl_3$ mixtures. This could explain why no phase separation was observed with sodium molybdate. **c,** Microscopic pictures of the RNA solutions supplemented with 10 mM salts. These pictures indicate that the RNA phase has different properties depending on the salt used. Although RNA with $MgCl_2$, $CaCl_2$, $CoCl_2$, and $ZnCl_2$ formed condensates, RNA with $MnSO_4$, $CuCl_2$, $FeSO_4$, and $FeCl_3$ formed condensates and bigger aggregates (or precipitates). The formation of aggregates correlates with the observation of distinct pellets in panel **a**, while condensates formed a thin, barely visible layer on the side of the tube in panel **a**. Scale bars correspond to 50 µm.

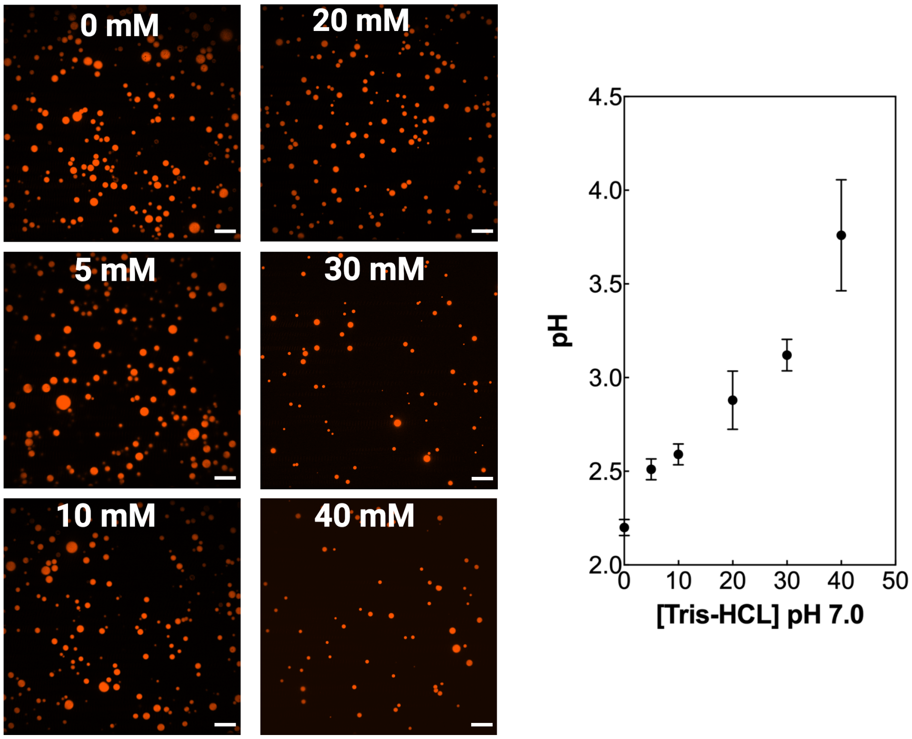


Supplementary Figure S 3: **Microscopic images of the RNA condensates obtained at different concentrations of Tris-HCl buffer pH 7.0 and graph reporting the pH of the entire solution measured with a pH meter**. RNA was used at 10 g/L and the phase separation was triggered using (100 mM) $MgCl_2$. Scale bars correspond to 20 µm.

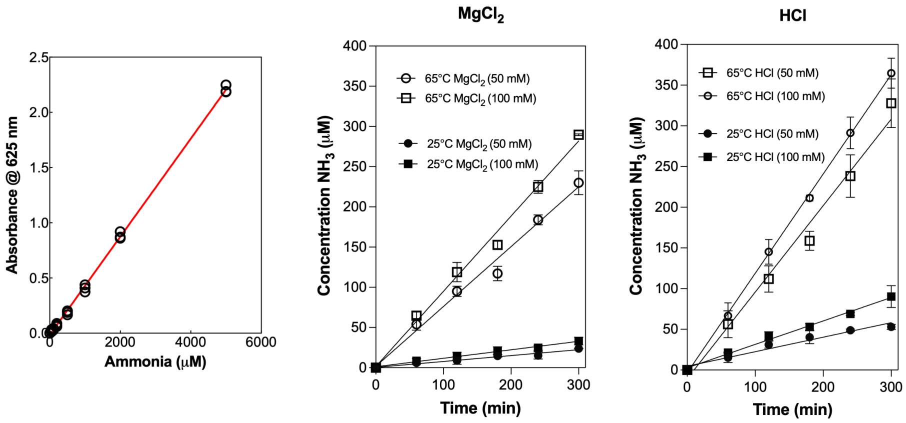


Supplementary Figure S 4: **RNA deamination analysis under phase separation conditions. Left panel:** Standard curve for ammonia detection at 625 nm. The linear relationship ($R^2 > 0.99$) was used to convert absorbance values to $NH_3$ concentrations. **Middle panel:** Time course of RNA deamination with $MgCl_2$ at different concentrations and temperatures. Open symbols represent 65°C conditions, filled symbols represent 25°C conditions. Circle symbols: 50 mM $MgCl_2$; square symbols: 100 mM $MgCl_2$. **Right panel:** Time course of RNA deamination with HCl at different concentrations and temperatures. Open symbols represent 65°C conditions, filled symbols represent 25°C conditions. Circle symbols: 50 mM HCl; square symbols: 100 mM HCl. Experimental conditions used in the experiment. RNA concentration: 7.5 g/L (comparable to phase separation conditions). $NH_3$ release was measured using the Berthelot reaction. Data points represent mean ± standard deviation of duplicate measurements. At 25°C, minimal deamination occurs over 4-8 hour timeframes typical of phase separation experiments, confirming RNA integrity is maintained under standard experimental conditions. Therefore, under our standard experimental conditions (25°C, 3-4 hour timeframes), RNA maintains excellent structural integrity with phase separation behavior unaffected by the minimal deamination observed.

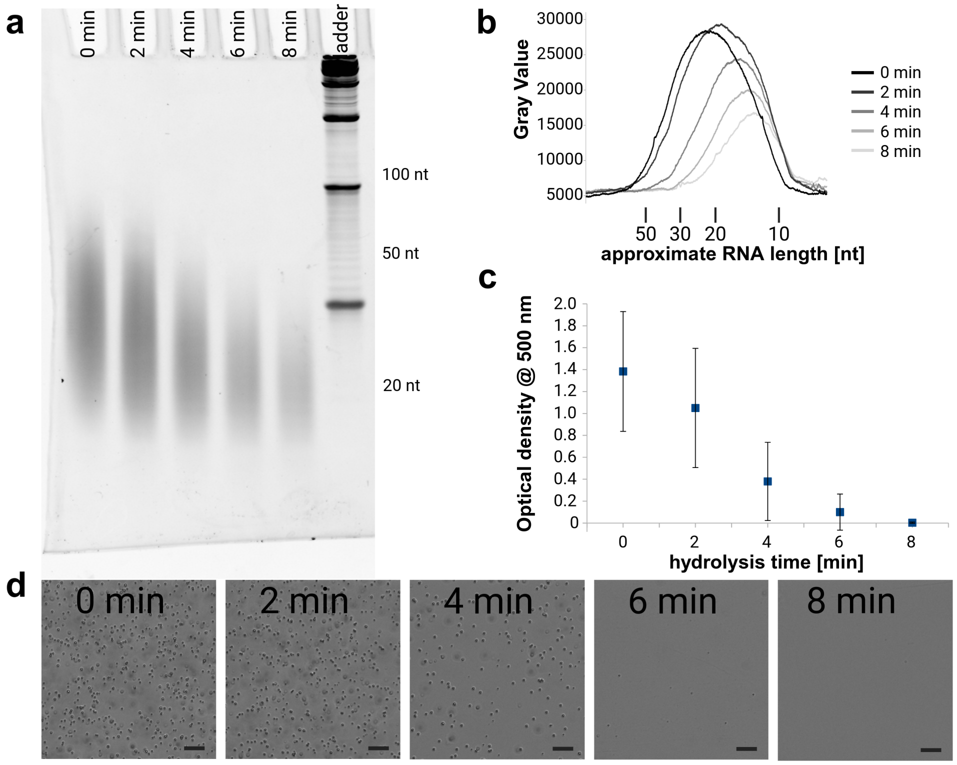


Supplementary Figure S 5: **RNA degradation by alkaline hydrolysis prevents phase separation.** **a,** Representative PAGE-gel showing the RNA size distribution before and after hydrolysis. An RNA solution of 3 g/L was incubated with 50 mM sodium hydroxide at 75°C for the indicated time. Afterwards, the solution was neutralized by adding 55 mM HCl. 1 µL of the treated RNA solution was loaded on the gel. **b,** The size distribution of the hydrolyzed RNA as estimated from the gel (shown in panel a) using ImageJ plotting. The size was estimated using the DynaMarker RNA Low II or FAM-labeled ribozyme substrate (see SI Table 2) hydrolyzed for 3 minutes under the conditions described for panel a. **c,** Optical density of phase separated RNA after alkaline hydrolysis. Phase separation was triggered by adding 50 mM $MgCl_2$. Values presented are averages and standard deviations for 4 experiments. **d**, Microscopic pictures show RNA condensates forming from RNA hydrolyzed for up to 4 - 6 minutes. Alkaline hydrolysis of around 6 - 8 minutes prevents RNA condensation. Scale bars are 20 µm.

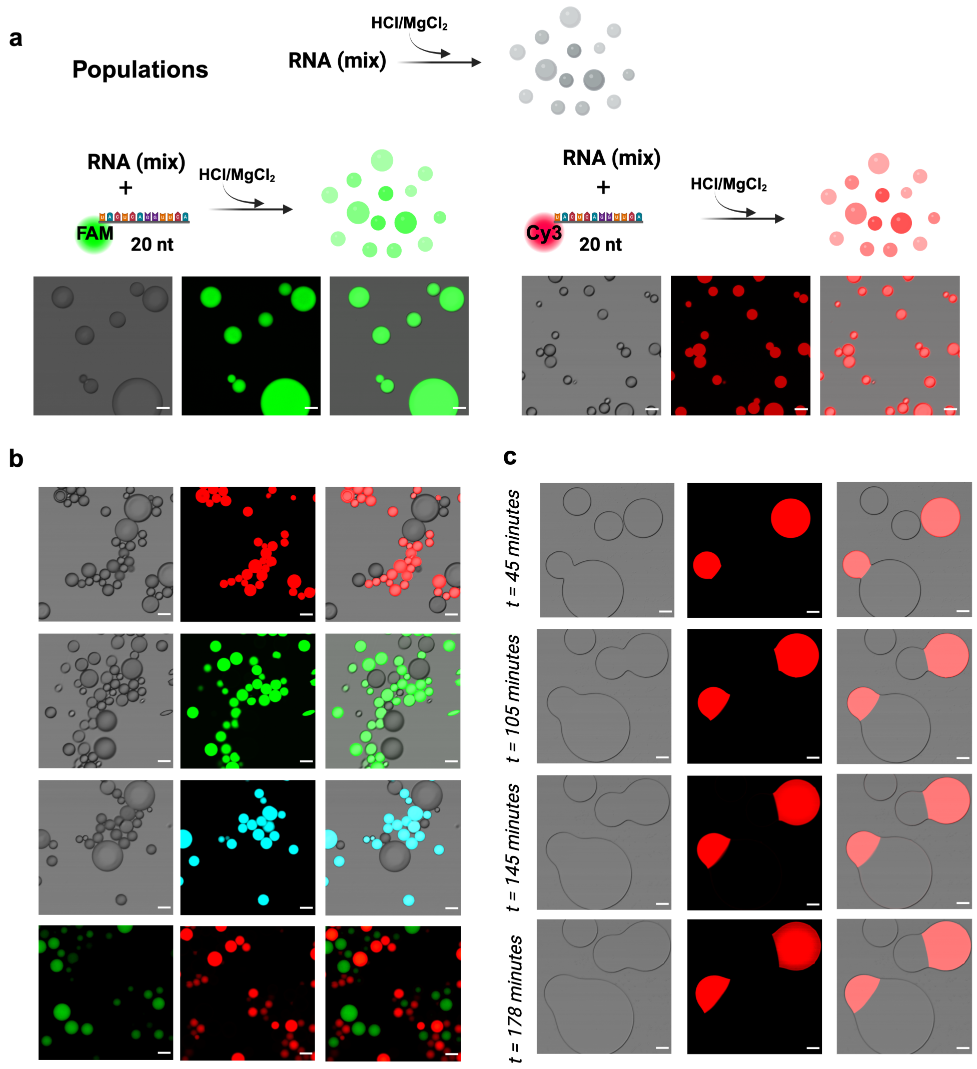


Supplementary Figure S 6: **Experiments using different RNA condensates population. a,** Schema illustrating the preparation of different ssRNA and ssDNA droplet populations. Fluorescent RNA droplet populations were prepared by compartmentalization of 20 nt ssDNA and ssRNA labeled with a fluorophore (FAM, Cy3 or Cy5, to exclude that the probe could drive nucleotide migration). To evaluate the transfer of the fluorescent cargo we used 2.5 µM as final concentration of the 20 nt labeled nucleic acids. **b,** The droplet populations kept their identities for prolonged periods of time after mixing; no transfer of the fluorescent cargo to unlabeled condensates was observed. It is interesting to note how many condensates form grape-like structures without merging. **c,** Fusion events over time between two different populations: one population contains a 20 nt Cy3 labeled RNA and the other population is without a probe. During fusion events the identities were preserved by the formation of sharp boundaries in the resulting bigger droplet. The time of the experiment was around 3 hours and limited by the wetting of the RNA condensates on the glass slide. The RNA condensate populations have been prepared by using 10 g/L RNA in the presence of 100 mM $MgCl_2$ or HCl. Note that we have mixed populations prepared using or HCl or $MgCl_2$. Scale bars correspond to 25 µm.

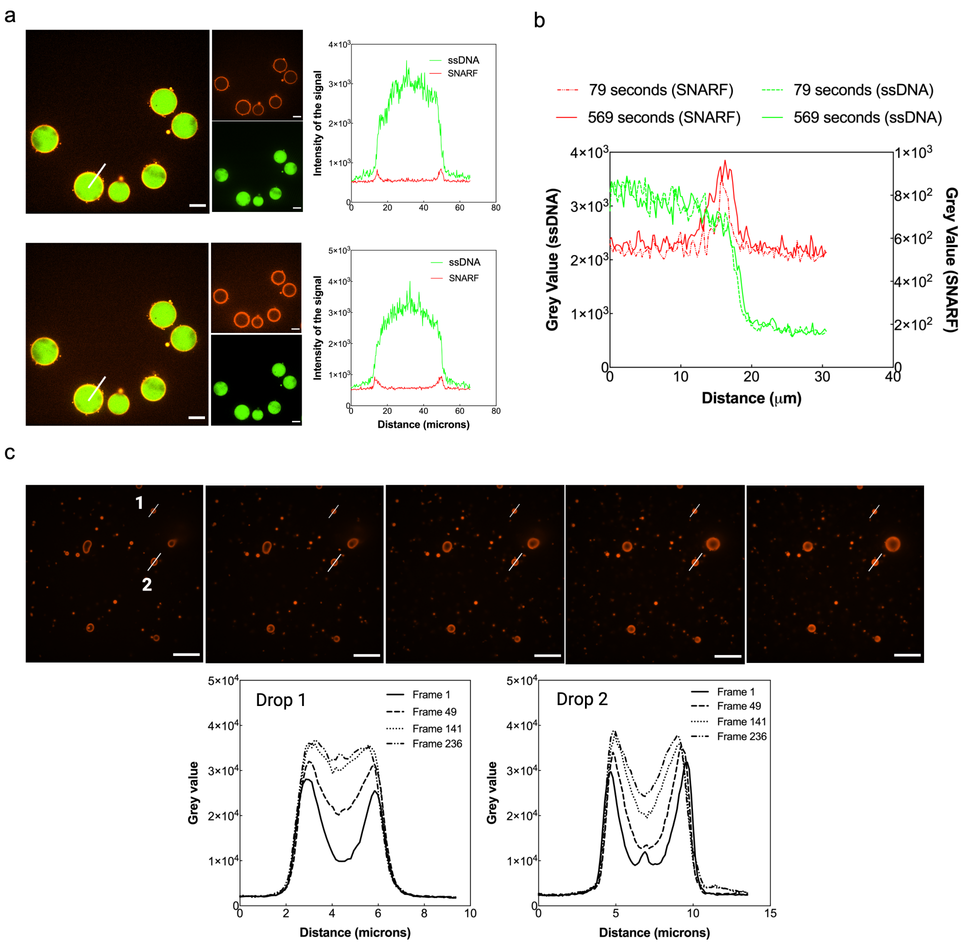


Supplementary Figure S 7: **Diffusion of small molecules within the RNA condensates**. The RNA condensates have been prepared using 10 g/L RNA and 100 mM $MgCl_2$. SNARF™-4F 5-(and-6)-carboxylic acid fluorescent probe was added at a final concentration of 5-10 µM, Rhodamine 6G at 20 µM. **a,** SNARF™-4F 5-(and-6)-carboxylic acid diffusion (red) at 79 (top) and 569 seconds (bottom) within pre-formed condensates containing FAM-labeled DNA (green; FAM-CACGATCtagttgagctGTCTACGcatgCGTAGACgttgaagGATCGTGatatata). **b,** Analysis of the intensity of the signal reported as grey value at two different time points. The red signal refers to SNARF while green one to FAM-labeled DNA partitioned in the condensates. **c,** Analysis of the diffusion of Rhodamine 6G within two condensates of different size at several frames. Scale bars correspond to 25 µm.

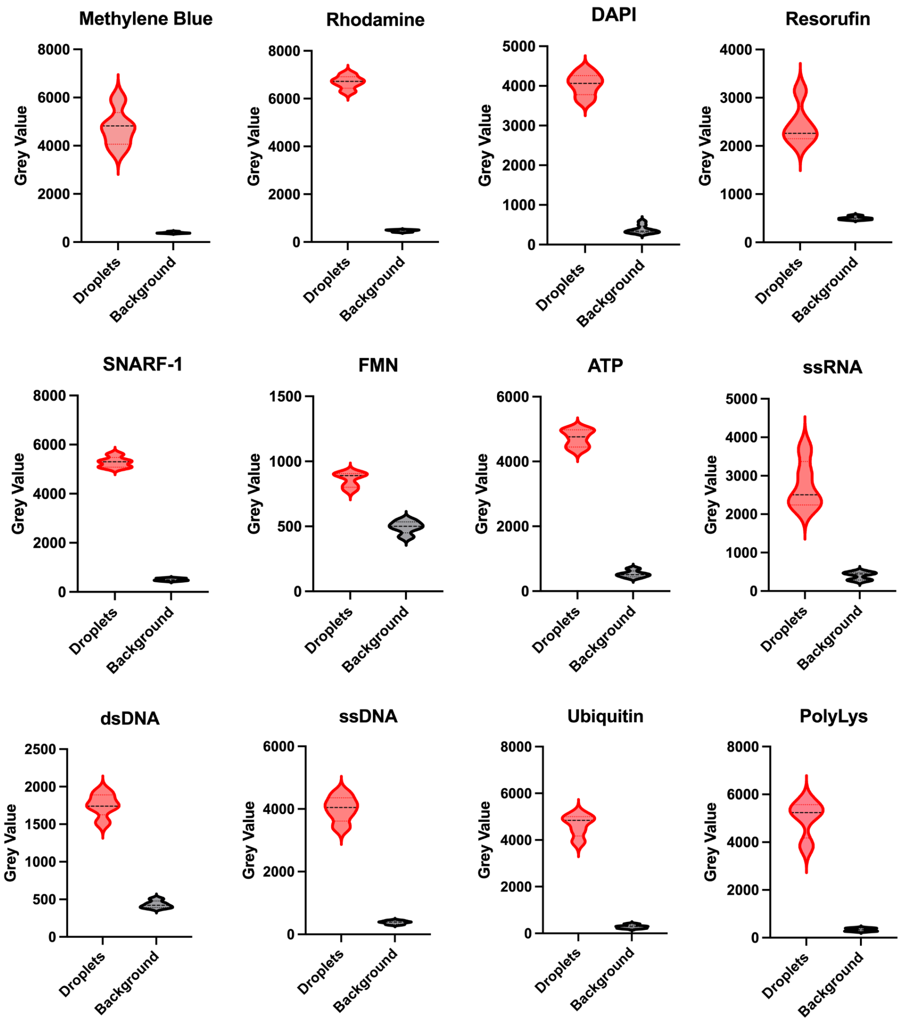


Supplementary Figure S 8: **Intensity reported as grey value of the organic molecules and peptides within and outside the RNA condensates**. The molecules have been added before triggering phase separation at different concentrations as reported in the main manuscript. The analysis has been performed using ImageJ and more than 10 condensates for each molecule have been analyzed from different experiments. The RNA phase separation has been triggered using both $MgCl_2$ or HCl at different concentration (from 10 mM to 100 mM) in the presence of 10 g/L RNA.

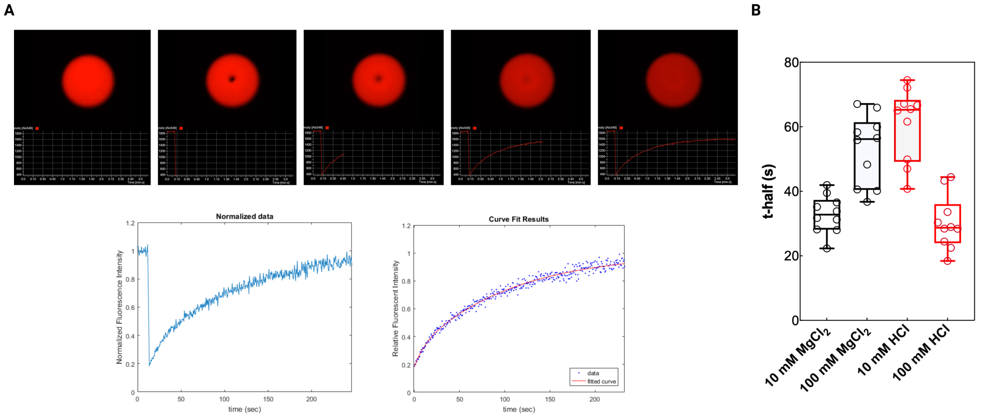


Supplementary Figure S 9: **FRAP analysis demonstrates liquid-like behavior of RNA condensates across different ionic conditions. A,** Representative FRAP experiment showing fluorescence recovery after photobleaching of rhodamine B-labeled RNA droplets (100 mM $MgCl_2$). Top panel: Sequential microscopy images captured at different time points during the recovery process, showing the photobleached spot (dark region) and subsequent fluorescence recovery. The images display both the fluorescence channel (upper portion) and corresponding intensity profile analysis (lower grid portion). Bottom panels: Quantitative analysis showing normalized fluorescence intensity recovery over time (left graph) and corresponding exponential curve fitting results (right graph, with experimental data points in purple and fitted curve in red). **B,** Quantitative comparison of fluorescence recovery half-times (t-half) measured across the ionic conditions used throughout this study: 10 mM $MgCl_2$ (black), 100 mM $MgCl_2$ (black), 10 mM HCl (red), and 100 mM HCl (red). Box plots display median values, interquartile ranges, and individual data points. RNA condensates were formed using 7.5 g/L total yeast RNA with different concentrations of $MgCl_2$ and HCl (10 and 100 mM) to trigger phase separation. Rhodamine 6G (R6G, 35 µM) was used as a fluorescent tracer for FRAP analysis. Data represent analysis of $n = 10$ droplets per condition.

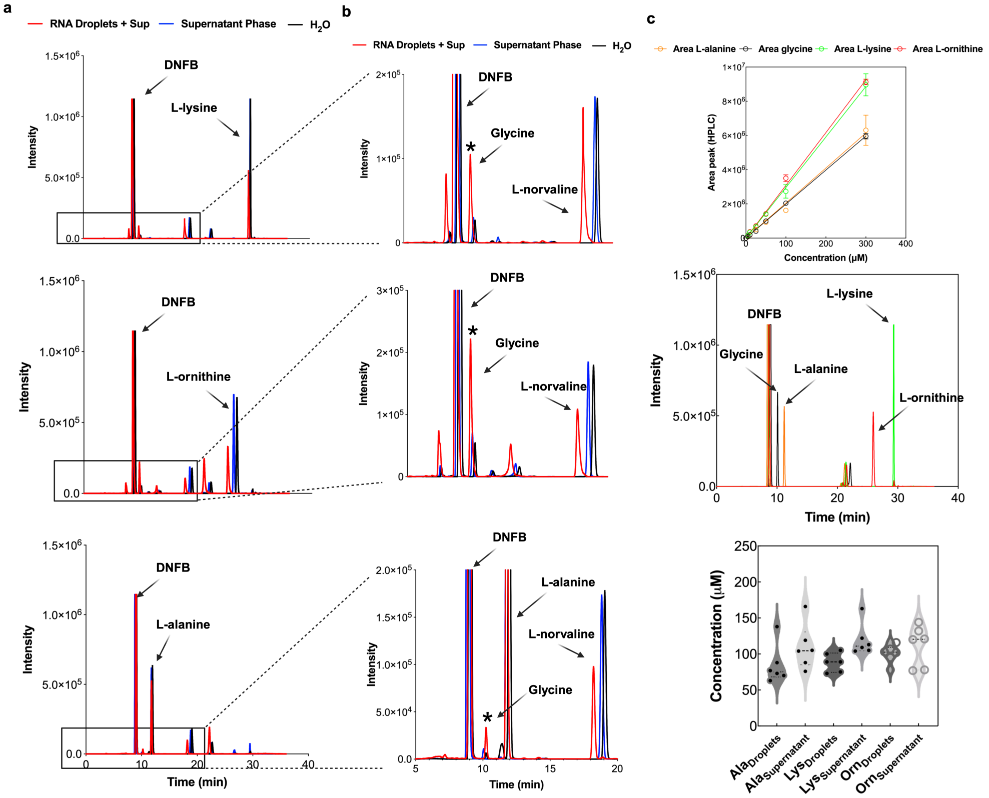


Supplementary Figure S 10: **HPLC chromatographic profiles, calibration curves and amino acids partitioning within the phases related to the experiments performed for glycine determination**. **a,** Chromatographic profiles recorded after derivatization of the amino acids using 1-Fluoro-2,4-dinitrobenzene (DNFB). Top to bottom: HPLC chromatogram of the experiments performed using l-alanine, l-ornithine and l-lysine (10 mM) as amino donor for the production of glycine in the presence of glyoxylate (20 mM). The red line corresponds to the chromatographic profile of the RNA condensates, while blue and black lines report the profiles obtained in the supernatant phase and water, respectively. **b,** Zoom-in of the chromatographic profiles reported in panel **a** which highlight the peak of the glycine with a retention time of 10 mL after that of the derivatizing agent DNFB. The l-norvaline has been used as internal control for the derivation process. **c,** Calibration curves of the amino acids in the two phases used in the experiments retrieved by calculating the area of the HPLC peaks at different amino acids concentrations and distribution of the amino acids between the two phases used at a final concentration of 10 mM. The experiments have been performed using 5 g/L RNA in the presence of 100 mM $MgCl_2$. All the components have been added just before triggering phase separation.

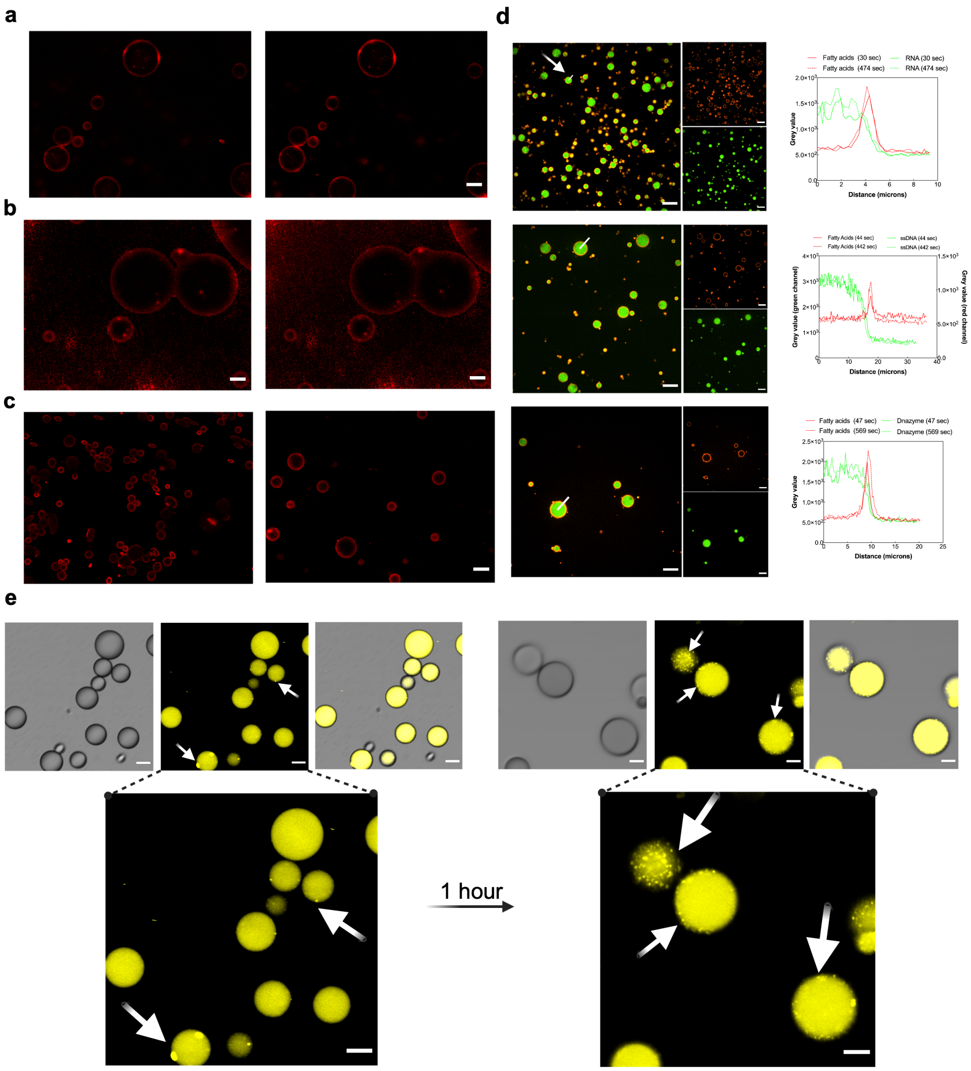


Supplementary Figure S 11: **Localization of fatty acids and phospholipids added to pre-formed RNA condensates**. **a,** Localization of undecylic acid (or C11 undecanoic acid) and **b,** Localization of lauric acid (or C12 dodecanoic acid) on the edges of the RNA condensates after 10 minutes (left panel) and 80 minutes (right panel). Both fatty acids have been used at a final concentrations of 5 µM. During the experimental time, no signal has been measured within the RNA condensates. **c,** Localization of the fatty acids (C11 and C12) on the edges of small condensates after 60 minutes (5 µM). **d,** Analysis of the signal intensity of the labeled fatty acids around the RNA condensates over time. The RNA condensates contain FAM labeled RNA 2.5 µM. The analysis of the intensity indicates that the fatty acids localize to the edges of the condensates, without entering in the condensates. **e,** Partitioning of phospholipid ($\beta$-BODIPY C12-HPC, Hexadecanoyl-sn-Glycero-3-Phosphocholine, used at a final concentration of 5 µM) within the RNA condensed condensates. Fatty acids and the phospholipid have been added after condensates formation, using both $MgCl_2$ or HCl (both at 100 mM). Scale bars correspond to 25 µm.

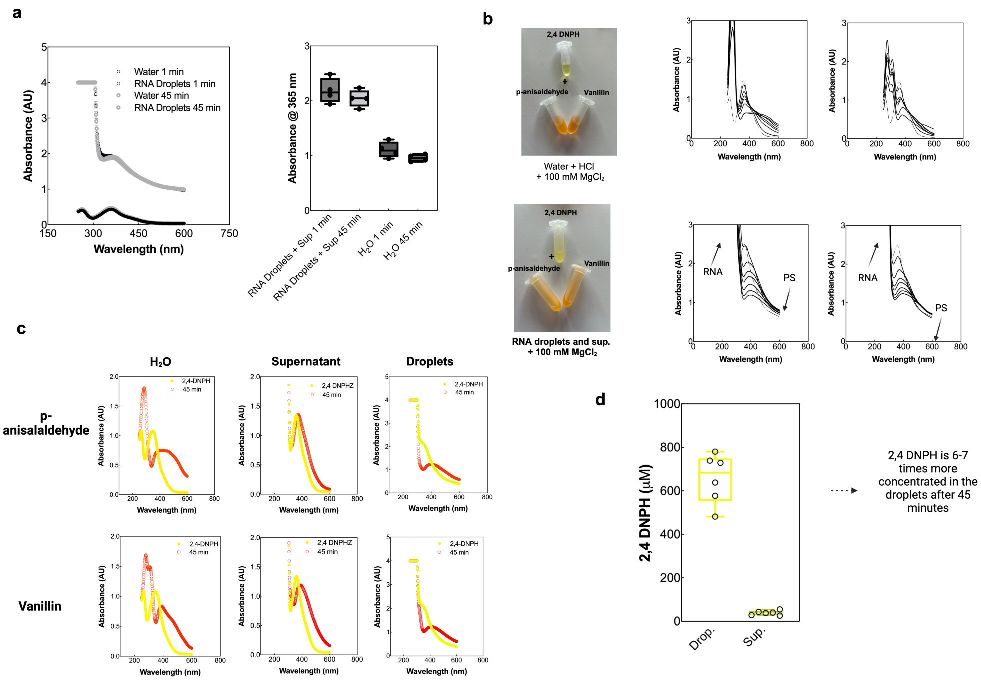


Supplementary Figure S 12: **Experimental set-up of the addition reaction (C-N bond formation) and absorbance data within the condensates. a,** Absorbance over time of the hydrazinic compound 2,4-dinitrophenylhydrazine (2,4-DNPH) used at 200 µM final concentration in water and RNA condensates under acidic conditions. 2,4-DNPH interacts with protein and nucleic acids. Therefore, we check its signal stability over time before recording any experimental data. After 45 minutes under agitation, the signal of 2,4-DNPH in condensates was stable as compared to water. **b,** Addition reaction within RNA condensates compared to dilute conditions. Image and absorbance spectra of hydrazone formation in water and RNA condensates using 2,4-DNPH with p-anisaldehyde and vanillin over time. 2,4-DNPH has been used at a final concentration of 200 µM and the aldehydes at a final concentration of 400 µM. The aldehydes were added along with 100 mM $MgCl_2$ which triggers phase separation. Differences in the absorbance maximum between the diluted conditions (top graphs) and in the presence of RNA condensates (bottom graphs, marked with arrows indicating RNA and PS). The reactions performed in acidic water (top), show the formation of the product (around 400-450 nm) while with RNA condensates (bottom) the signal is covered by the signal of the phase separation (turbid solution). The signal of the 2,4-DNPH around 300 nm is also completely covered by the absorption of RNA. **c,** Absorbance spectra of the reaction between 2,4-DNPH and p-anisaldehyde and vanillin in water, supernatant phase, and RNA condensates. The 2,4-DNPH spectra is represented as yellow dots while in orange empty dots report the spectra 45 minutes after aldehyde addition. The color of the solutions turn from yellow to bright orange (p-anisaldehyde) or to dark orange (vanillin) after 1 hour incubation in acidic conditions due to the formation of (precipitated) hydrazone. **d,** Partitioning of the 2,4-dinitrophenylhydrazine (yellow color) between the condensates and supernatant phase used at (150 µM) after one hour incubation. The condensates were centrifuged and the supernatant removed. Then, the droplet phase was resuspended at pH 7.4 in potassium phosphate buffer 200 mM and read at the spectrophotometer using $\varepsilon_{365} = 22000$ $M^{-1}cm^{-1}$.[24]

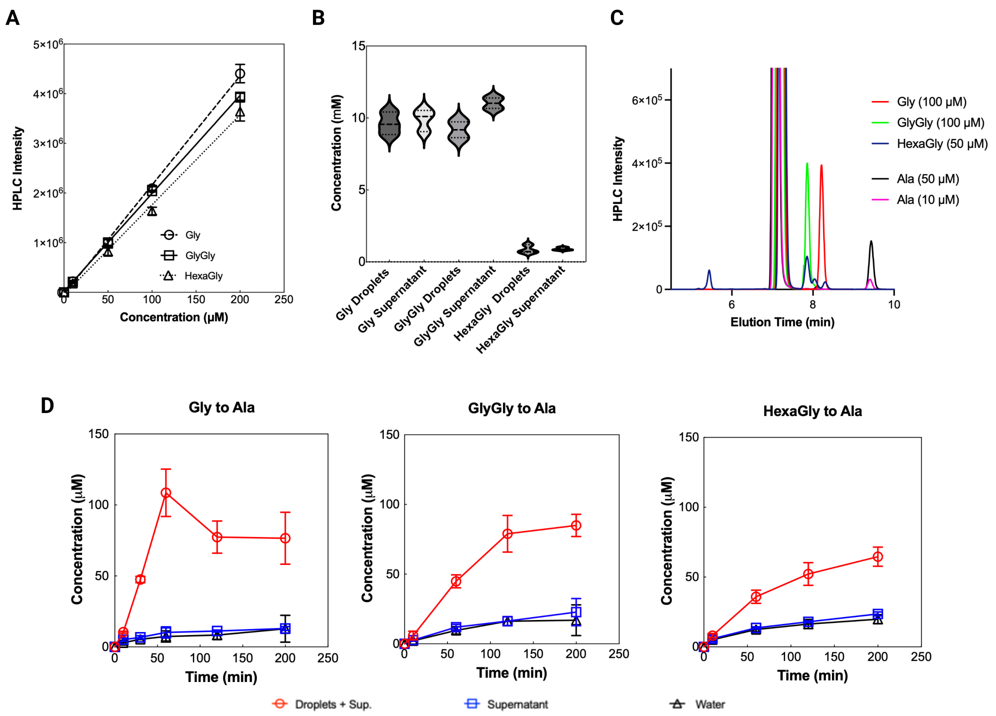


Supplementary Figure S 13: **Enhanced amino acid transamination in RNA condensates using pyruvate as $\alpha$-ketoacid.** **A,** Calibration curves for glycine (Gly, circles), diglycine (GlyGly, squares), and hexaglycine (triangles) obtained by HPLC analysis at four different concentrations (0, 10, 50, 100, 200 µM). **B,** Partitioning analysis of amino acids and peptides between RNA condensate (droplets) and supernatant phases after centrifugation. Analytes were added at concentrations of 10 mM for Gly and GlyGly, and 1 mM for hexaglycine. **C,** Representative HPLC chromatograms and elution times of the analyzed compounds after derivatization with DNFB. **D,** Time-course analysis of transamination reactions converting glycine substrates to alanine in the presence of pyruvate. Reaction conditions: 10 mM pyruvate, 7.5 mM Gly and GlyGly, 1 mM hexaglycine, in RNA condensates (red circles), supernatant phase (blue squares), or water control (black triangles). Aliquots were derivatized with DNFB and analyzed by HPLC. RNA condensates demonstrate significant enhancement of alanine formation with reaction rates enhanced approximately 4 - 10 fold over water controls.

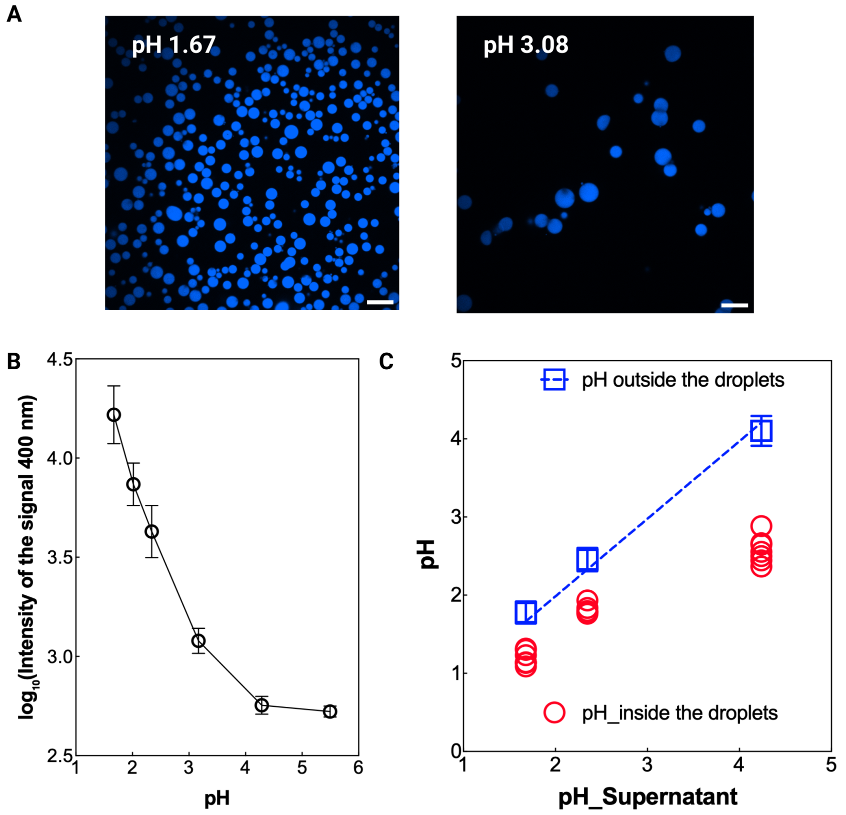


Supplementary Figure S 14: **LysoSensor-based measurement of pH within RNA condensates.** **A,** Representative fluorescence microscopy images of RNA droplets containing LysoSensor at external pH 1.67 (left) and pH 3.08 (right). Higher blue fluorescence intensity indicates lower internal pH. **B,** Calibration curve for LysoSensor pH probe. The probe exhibits pH-dependent fluorescence intensity, with higher intensity at lower pH values. Log10 (fluorescence intensity at 400 nm) is plotted versus solution pH measured with a standard pH electrode. For the calibration curve, solutions of a certain pH (with and without dissolved RNA) were measured by microscopy to verify that RNA does not affect the probe response; no difference was observed. Each point represents the mean ± SD of duplicate measurements. **C,** Quantitative comparison of pH measured inside individual RNA droplets (red circles) versus external solution pH (blue squares). Each red circle represents a single droplet measurement; blue squares show external pH measured with a standard electrode. The dashed blue line indicates pH identity (internal pH = external pH). Systematic displacement of all measurements below the identity line demonstrates consistent acidification within RNA condensates, with internal pH ranging from 1.3 to 2.9 across the tested conditions. No droplets showed internal pH equal to or higher than external pH, indicating universal acidification. RNA condensates were formed using 7.5 g/L total yeast RNA and 50 mM $MgCl_2$. External pH was varied using Tris-HCl buffer at different concentrations (0, 10, 25, 50, 75, 100 mM) corresponding to the pH values reported in the graph. LysoSensor pH probe (20 µM final concentration) was added after droplet formation. Scale bars are 20 µm.

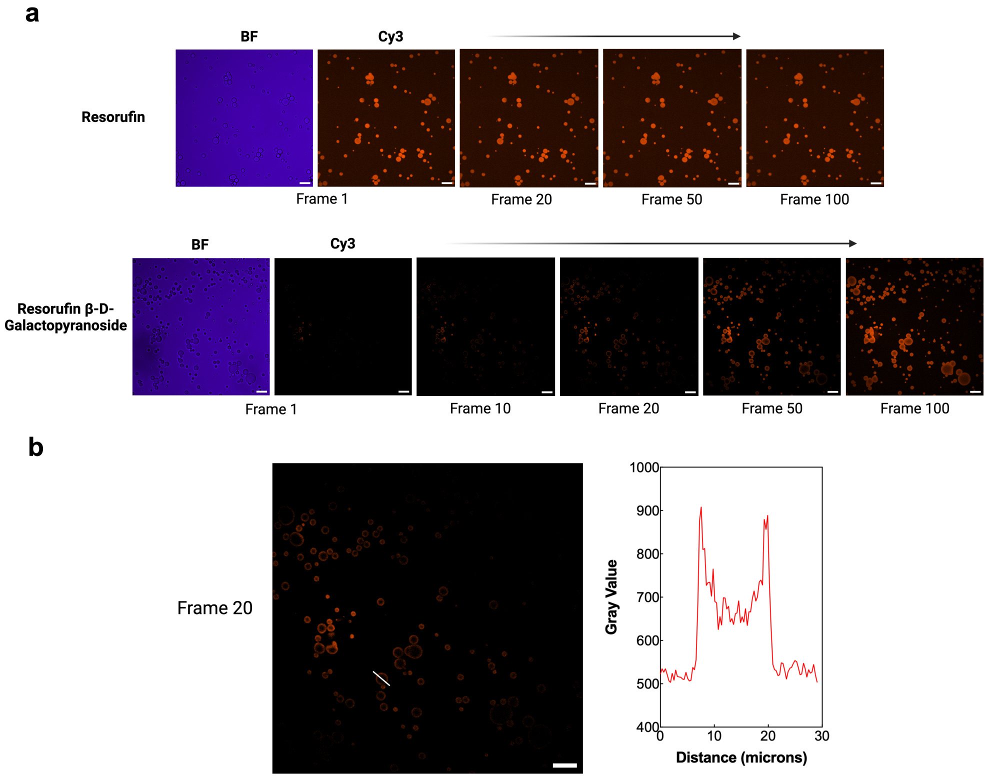


Supplementary Figure S 15: **Spatial dynamics of catalytic activity within RNA condensates.** Real-time fluorescence microscopy tracking hydrolysis of resorufin $\beta$-D-galactopyranoside (100 µM) within RNA condensates. The substrate is non-fluorescent; hydrolysis releases fluorescent resorufin. **a,** The top panel shows a control experiment with pre-formed resorufin (100 µM) displaying immediate, uniform fluorescence distribution throughout the field. The bottom panel demonstrates the substrate hydrolysis reaction with edge-initiated catalytic activity. Brightfield (BF) images show condensate morphology while the Cy3 channel captures resorufin fluorescence (product of hydrolysis). Fluorescent product first appears at droplet peripheries (Frame 10 - 20) and progressively fills the condensate interior (Frames 50 - 100). **b,** Higher magnification view of Frame 20 showing the edge-to-center fluorescence gradient (left panel) with corresponding fluorescence intensity profile across a representative condensate (right panel, white arrow indicates line of measurement). This spatial progression indicates that substrate molecules experience an altered chemical environment immediately upon condensate entry, promoting glycosidic bond hydrolysis. The condensed phase retains and concentrates reaction products, and the acidified condensate environment enhances this hydrolysis reaction. The edge-to-center progression suggests condensates function as spatially organized reaction vessels with distinct microenvironments. RNA condensates were formed using 7.5 g/L total yeast RNA in the presence of 100 mM $MgCl_2$ (similar results observed with 25 and 50 mM $MgCl_2$). Scale bars are 20 µm.

**a**

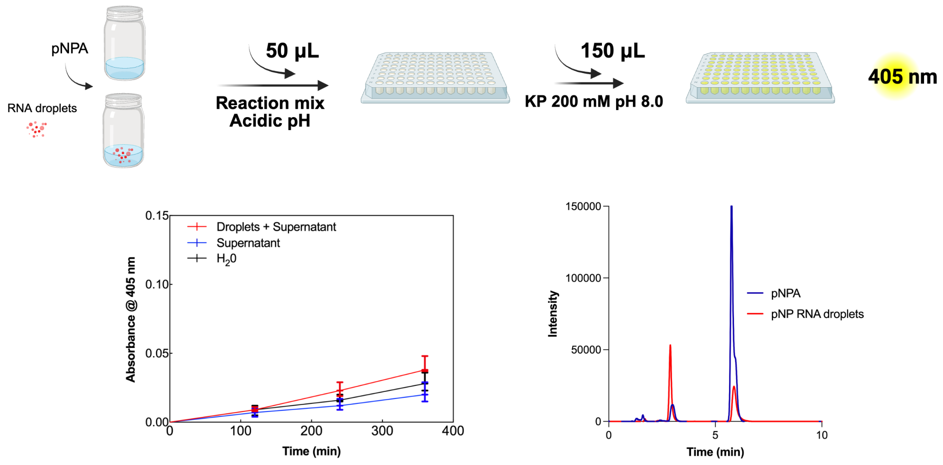


**b**

| Condition | 1 mM pNPA (M/s) | 4 mM pNPA (M/s) |
|---|---|---|
| RNA Droplets | $0.8 \pm 0.3 \times 10^{-9}$ | $3.0 \pm 0.7 \times 10^{-9}$ |
| Diluted Phase | $0.4 \pm 0.1 \times 10^{-9}$ | $1.9 \pm 0.4 \times 10^{-9}$ |
| Water | $0.7 \pm 0.3 \times 10^{-9}$ | $2.6 \pm 0.3 \times 10^{-9}$ |

Supplementary Figure S 16: **Schema reporting the experiments performed for the pNPA hydrolysis. a,** The pNPA has been added along with $MgCl_2$ (100 mM) which triggers phase separation. Since the signal of the pNP formed under acidic conditions (around 320 nm) was covered by the high RNA concentration in the condensed samples, we decided to analyze the signal under basic pH condition (pH 8.0) (conversion from protonated to deprotonated form of the para-nitrophenolate anion). In detail, 50 µL of the reaction mix were mixed with 150 µL of potassium phosphate buffer (200 mM) pH 8.0. Under this condition, the pNP turns yellow (para-nitrophenolate anion) with maximum absorbance at 405 nm. Graphs reporting the pNPA hydrolysis under acidic condition obtained in the presence of RNA condensates, with the supernatant phase, and in water for 6 hours using pNPA concentration 1 mM, left panel). On the right, HPLC peaks of pNPA and pNP used to confirm the hydrolysis of pNPA to pNP and acetate in the samples. **b,** Tables reporting the kinetic analysis regarding the acid hydrolysis of pNPA using a zero-order model (see detailed explanation in the SI section **Hydrolysis of pNPA under acidic conditions**”).

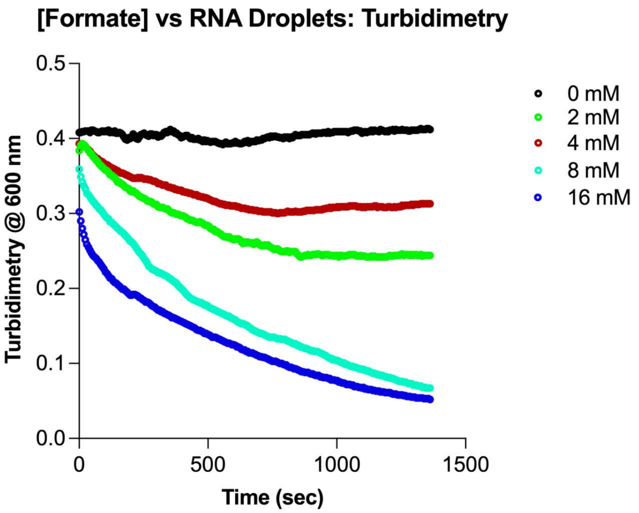


Supplementary Figure S 17: **Formate effect on RNA condensates.** Turbidimetry experiments related to the RNA condensates in presence of different concentrations of formate added to the condensates. The change of the pH of the overall solution due to the formate addition dissolve the condensates.

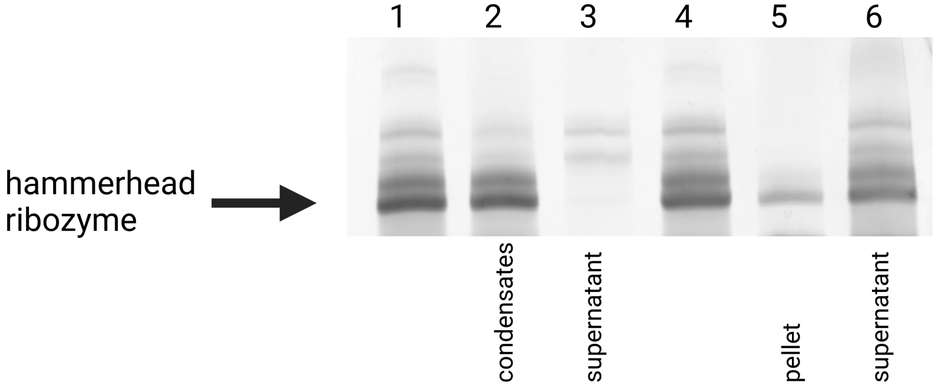


Supplementary Figure S 18: **Partitioning of the hammerhead ribozyme between condensates and supernatant.** Sybr Gold stained denaturing PAGE gel showing 125 ng hammerhead ribozyme (46 nt: GGGCGAGGUACAUCCAGCUGACGAGUCCCAAAUAG-GACGAAAUGCC) in the presence of 100 mM $MgCl_2$, 10 mM KCl, 5% propylene glycol, and 10 mM bis-tris-propane (pH 6.2 for lane 1, 2, 3 and pH 7.2 for lane 4, 5, 6) in a total of 25 µL. Lanes 2, 3, 5, 6 also contain 4 g/L yeast RNA. The samples were incubated at 37 °C for 45 minutes, spun down at 21,000 g for 2 minutes to separate condensates and supernatant. Condensates were resuspended in 25 µL water and all samples were then precipitated with ethanol and sodium acetate, before loading on the gel. Phase separation occurred in the samples shown in lane 2 and 3 (low pH, yeast RNA, and magnesium present). There, the hammerhead ribozyme is found exclusively inside the condensates. In lanes 5 and 6 no phase separation occurred due to the higher pH. There, the ribozyme is mostly found in the supernatant. Ribozyme in the pellet (lane 5) was probably sticking to the tube walls with residual liquid when removing the supernatant or could be inside few and small condensates still present.

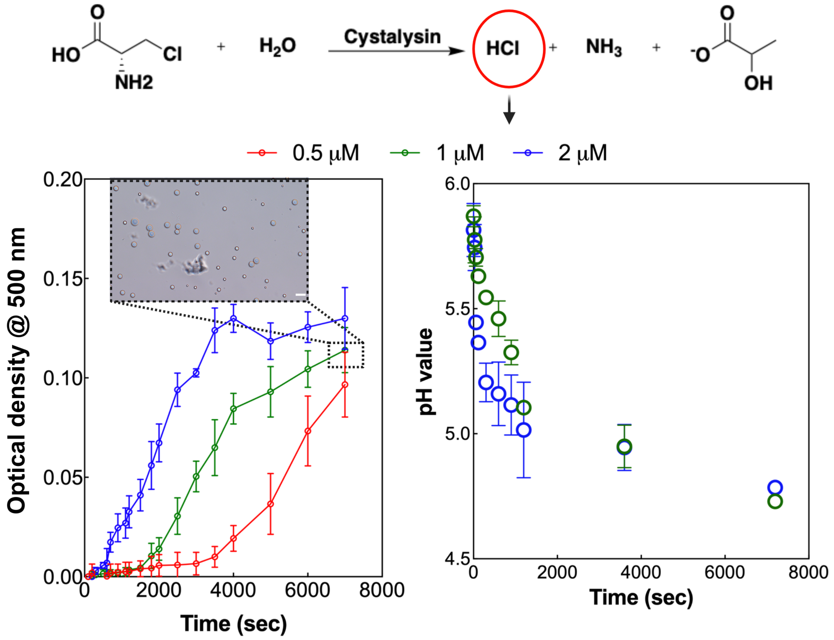


Supplementary Figure S 19: **RNA phase separation is triggered by enzymatic activity.** Schema of the catalytic reaction ($\alpha, \beta$ elimination) performed by cystalysin using $\beta$-chloroalanine as substrate. Graph reporting the increase of the optical density signal monitored at 500 nm in the presence of active cystalysin. The signal is directly related to RNA condensates formation mediated by the released HCl which is a product of the cystalysin reaction. The PLP dependent enzyme cystalysin has been used at different concentrations (0.5 µM red dots and line, 1 µM green dots and line, and 2 µM blue dots and line) in the presence of 375 ng/µL RNA, exogenous PLP 20 µM, and (1 mM) $\beta$-chloroalanine. The enzyme produces HCl that decreases the pH of the solution thereby triggering RNA phase separation as demonstrated by the increasing values of optical density at 500 nm as well as by the bright-field microscope image captured after 3 hours of reaction. Scale bars correspond to 25 µm.

**a**

**Kinetic analysis (p-anisaldehyde, 2,4-dinitrophenylhydrazone)**

| Experimental Condition | k ($\mu M^{-1} min^{-1}$) | $t_{1/2}$ (min) |
|---|---|---|
| **RNA Droplets + sup.** | $(4.5 \pm 0.6) \times 10^{-4}$ | $2.7 \pm 0.4$ |
| **Diluted phase** | $(5.0 \pm 0.5) \times 10^{-5}$ | $24 \pm 2$ |
| **Water** | $(1.9 \pm 0.1) \times 10^{-4}$ | $6.7 \pm 0.6$ |

**Kinetic analysis (vanillin, 2,4-dinitrophenylhydrazone)**

| Experimental Condition | k ($\mu M^{-1} min^{-1}$) | $t_{1/2}$ (min) |
|---|---|---|
| **RNA Droplets + sup** | $(1.7 \pm 0.3) \times 10^{-4}$ | $7 \pm 1$ |
| **Diluted phase** | $(2.6 \pm 0.3) \times 10^{-5}$ | $47 \pm 6$ |
| **Water** | $(1.0 \pm 0.1) \times 10^{-4}$ | $12 \pm 1$ |

| Experimental Condition | p-Anisaldehyde $t_{1/2}$ (min) | Vanillin $t_{1/2}$ (min) | Acceleration Factor (p-Anisaldehyde)* | Acceleration Factor (Vanillin)* |
|---|---|---|---|---|
| **Diluted phase** | $24 \pm 2$ | $47 \pm 6$ | 1.00* | 1.00* |
| **RNA Droplets + sup.** | $2.7 \pm 0.4$ | $7 \pm 1$ | 8.8 | 6.6 |
| **Water** | $6.7 \pm 0.6$ | $12 \pm 1$ | 3.7 | 3.8 |

**b**

**Transamination reaction**

| Amino Acid | Condition | Rate (μM/min) | Max Yield (μM) |
|---|---|---|---|
| L-Alanine | RNA Droplets | 14 | 493 |
| | Diluted Phase | 4 | 137 |
| | Water | 1 | 57 |
| L-Lysine | RNA Droplets | 38 | 1484 |
| | Diluted Phase | 7 | 462 |
| | Water | 2 | 89 |
| L-Ornithine | RNA Droplets | 107 | 4192 |
| | Diluted Phase | 14 | 995 |
| | Water | 10 | 483 |

Supplementary Figure S 20: **Tables reporting the kinetic analysis regarding the experiments related to hydrazones and glycine formation. a,** Table reporting the rate and half-life values obtained using the second order kinetic analysis for the formation of the hydrazones. Table reporting the analysis of the half-life obtained under different experimental conditions used to calculate the acceleration factor. * $AccelerationFactor = t_{1/2}(dilutephase)/t_{1/2}(condition)$ - This represents how many times faster the reaction reaches 50% conversion compared to the diluted phase. (see supporting information **Addition reaction under acidic conditions** for more information). **b,** Table reporting the rates and the yield obtained during the transaminations experiments for all the amino acids used under three different conditions.
